# Integrative Multiomics Analysis of the Angiogenesis–EMT Interface in Head and Neck Squamous Cell Carcinoma

**DOI:** 10.64898/2026.11.05.764737

**Authors:** Sreyashi Das, Elimathi Natarajan, Rashmi Singh, Rizwan Shaikh

## Abstract

Angiogenesis and epithelial–mesenchymal transition (EMT) are key hallmarks that contribute to the progression of oral squamous cell carcinoma (OSCC), but whether the two programs vary simultaneously across tumors and how much of this reflects tumor composition remain unclear. We scored predefined, non-overlapping 20-gene angiogenesis and EMT signatures with a rank-based method in 342 primary oral cavity tumors from The Cancer Genome Atlas (TCGA-OSCC). The two programs were strongly correlated (Spearman ρ = 0.809, P = 2.7 × 10^−80^), and 2,396 bridge genes were associated with both programs across the transcriptome. The bridge genes were enriched in pathways associated with focal adhesion and extracellular matrix– receptor interaction. In the cellular case, fibroblast and endothelial marker scores were prominent and correlated with the program scores. The coupling was attenuated but remained positive after joint adjustment for tumor purity and stromal and immune scores (partial ρ = 0.406) and with an epithelial-restricted EMT score (ρ = 0.710). Further correlation was performed with a partial EMT (pEMT), which shows a significant positive correlation (ρ = 0.59) and an inverse correlation with tumor purity (ρ = −0.70). 146 final gene candidates were found using multi-omic evidence and candidate-level testing on the external cohort GSE30784; after adjusting for inferred cell fractions, 101 of these candidates supported the dual program. For thorough characterization, three candidates with an emphasis on OSMR, MICAL2, and LHFPL2 were chosen. Additional CPTAC data were used to confirm protein-level support, which was strongest for MICAL2. No significant connection was seen between the Interface score and overall survival (hazard ratio per 1-SD increase, 0.92; 95% CI, 0.78–1.08). These results describe a coordinated angiogenesis–EMT transcriptional state in OSCC that is attenuated but retained after accounting for tumor composition, and nominate candidate genes for experimental study.

## Introduction

Oral squamous cell carcinoma (OSCC) arises from the mucosal epithelium of the oral cavity, including the pharynx and larynx (Johnson et al., 2020), and accounted for an estimated 452,205 cases and 194,108 deaths worldwide in 2024 (IARC Global Cancer Observatory, 2024 estimates). These tumors are most often diagnosed at a later stage (Johnson et al., 2020), with survival decreasing as the disease spreads. Relative survival for the oral cavity and pharynx is 88.7% for localized disease, 69.7% for locally advanced disease, and 36.0% after distant metastasis (SEER, 2016–2022 data). These stages of the disease only report tumor metastasis, but not the biological programs that vary between tumors; these tumors are molecularly heterogeneous (The Cancer Genome Atlas Network, 2015).

All cancer growth and progression are linked to two major processes: new blood vessel formation (angiogenesis) and epithelial–mesenchymal transition (EMT), in which epithelial cells acquire migratory traits and become mesenchymal-type (Hanahan & Weinberg, 2011; Yang et al., 2020). These two processes are interlinked (Pomella et al., 2024). There is an interaction between elevated expression of E-cadherin and VEGF and overall survival rate in OSCC reported by prior meta-analysis (Zhao et al., 2013; Lorenzo-Pouso et al., 2023), where EMT is an intermediate state that drives metastasis; angiogenesis is particularly crucial for tumor growth and nourishment (Hanahan & Weinberg, 2011) and cannot be diagnosed by a single or few markers (Yang et al., 2020; Pal et al., 2021). In HNSCC, single-cell sequencing reported that the partial EMT (pEMT) program occurs at the leading edge of tumors, adjacent to cancer-associated fibroblasts (CAFs), and is associated with nodal metastasis (Puram et al., 2017); markers measured by immunohistochemistry were also associated with nodal metastasis (Parikh et al., 2019). Most transcriptomic studies in this field have reported these two hallmark programs separately as prognostic signature genes in OSCC (Ai et al., 2023; Chen et al., 2024).

Previous studies have associated angiogenic factors with both new vessel formation and EMT in OSCC (Pomella et al., 2024). Still, it remains underexplored whether angiogenesis and EMT programs differ simultaneously across patients’ tumors, and which genes or pathways are shared by both. This has been addressed by bulk tumor expression across many patients. Still, cautious handling is warranted, as bulk profiles contain a mixture of malignant cells, fibroblasts, endothelial cells, and immune cells. Additionally, differences in the proportion of cancer cells within tumor samples can influence correlations between gene expression levels across tumors (Aran et al., 2015). For instance, in colorectal cancer, mesenchymal expression signatures in bulk tumors mostly emerge from stromal cells (Isella et al., 2015). In HNSCC, the mesenchymal subtype, defined from bulk data, reflected fibroblast and myocyte content (Puram et al., 2017), and EMT signals in bulk tumors are difficult to distinguish from stromal signals (Tyler & Tirosh, 2021). A bulk EMT score is therefore not the same as EMT and pEMT in malignant cells, and an angiogenesis score that includes endothelial genes such as PECAM1, CDH5, and KDR partly reflects endothelial content.

These observations led us to investigate whether angiogenesis-and EMT-related transcriptional programs are coordinated across primary OSCC tumors and, if so, which specific genes and pathways accompany both programs, and how much of this coordination reflects tumor composition. We have hypothesized that tumors with elevated expression of angiogenesis programs would also show higher EMT expression, and that this coordination would extend beyond the signature genes to a broader set of genes associated with both programs. Because bulk tumors comprise both stromal and endothelial cells, we also tested whether the coordination reflects cellular composition rather than the state of malignant cells.

To investigate this, we primarily scored predefined, non-overlapping angiogenesis and EMT signatures in oral cavity tumors from TCGA-HNSC (The Cancer Genome Atlas Network, 2015). Further, we combined these into an angiogenesis–EMT coupling interface score. Additionally, we identified the bridge genes that are associated with both programs across the transcriptome and evaluated the contribution of tumor composition using tumor microenvironment (TME) marker scores, ESTIMATE purity (Yoshihara et al., 2013), single-cell RNA-seq data (GSE103322; Puram et al., 2017), cell-type deconvolution (Chu et al., 2022; Hu & Chikina, 2024), and a pEMT score, and we included multi-omic approaches such as copy-number, DNA methylation, and miRNA data as supporting evidence. To further rule out the data-dependent artifact, we assessed the program coupling in an external OSCC cohort (GSE30784; Chen et al., 2008), which also contributed to the selection of final candidates, and protein data in CPTAC-HNSCC (Huang et al., 2021), a mixed-site HNSCC comparison cohort rather than an independent OSCC validation cohort. OSMR, MICAL2, and LHFPL2 were three illustrative focused candidates selected for detailed characterization. Lastly, we evaluated the association between the Interface score and clinicopathological features and overall survival. Taken together, these report a coordinated transcriptional state and nominate candidate genes for future clinical validation.

## Materials and Methods

### Study cohort

All analyses used publicly available, de-identified data. TCGA head and neck squamous cell carcinoma (HNSC) RNA-seq data (The Cancer Genome Atlas Network, 2015; RSEM-normalized, log2(x + 1); TCGA_HNSC_HiSeqV2) and the HNSC clinical matrix were obtained from UCSC Xena (Goldman et al., 2020). Tumors were selected by keyword matching on the anatomic_neoplasm_subdivision field (oral cavity, tongue, floor of mouth, buccal mucosa, alveolar ridge, hard palate, lip and gingiva), keeping one primary tumor sample (type 01) per patient; because “tongue” also matched base of tongue, an oropharyngeal subsite, these tumors were included. The TCGA-OSCC cohort comprised 342 patients (oral tongue, 130; oral cavity not otherwise specified, 73; floor of mouth, 62; base of tongue, 27; buccal mucosa, 22; alveolar ridge, 18; hard palate, 7; lip, 3), selected without regard to HPV status (p16: 12 positive, 51 negative, 279 not recorded). The 20,213 of 20,501 genes with non-zero variance were used for gene-level tests. Gene-level GISTIC2 copy number (TCGA_HNSC_GISTIC2, UCSC Xena), Illumina HumanMethylation450 β-values and mature miRNA expression (miRNA_HiSeq_gene; log2(RPM + 1)), both from the UCSC Xena TCGA hub (version 2017-09-08), were matched to the same patients by barcode.

### External datasets

The gcRMA-normalized GSE30784 series matrix (GPL570; Chen et al., 2008) was downloaded from GEO; samples were classified from their GEO characteristics as OSCC (n = 167), normal oral mucosa (n = 45) or oral dysplasia (n = 17), and probes were collapsed to 22,829 gene symbols by their median. CPTAC-HNSCC gene-level RNA-seq (TPM), global proteomic and clinical data (Huang et al., 2021) were restricted to tumor profiles whose identifiers matched a clinical patient ID exactly, excluding 2 participants with normal tissue only, which gave 110 patients (oral cavity, 49; larynx, 49; oropharynx, 6; lip, 4; hypopharynx, 2). Protein values were log2 ratios, median-centered within each sample. The processed tables were obtained with the cptac Python package (version 1.5.14; Lindgren et al., 2021) on 13 September 2026, using its umich proteomics, broad RNA-seq and mssm clinical data. Processed GSE103322 single-cell data (Puram et al., 2017; log2(TPM/10 + 1); 23,686 genes) were used with the authors’ annotations: 2,215 malignant and 3,363 non-malignant cells were kept, 324 unclassified cells were excluded, 18 cells labeled “-Fibroblast” were merged with fibroblasts, and cells from primary tumors and lymph-node metastases (1,334 lymph-node cells) were both retained.

### Gene signatures and program scores

The angiogenesis signature comprised VEGFA, VEGFB, VEGFC, KDR, FLT1, FLT4, PDGFB, PDGFRB, ANGPT1, ANGPT2, TEK, PECAM1, CDH5, ENG, NRP1, NRP2, CXCL12, CXCR4, HIF1A and EPAS1, and the EMT signature comprised VIM, FN1, CDH2, SNAI1, SNAI2, TWIST1, TWIST2, ZEB1, ZEB2, MMP2, MMP9, MMP14, CDH1, OCLN, CLDN1, TJP1, TGFB1, TGFB2, TGFBR1 and TGFBR2. The genes were curated to represent VEGF ligands and receptors, PDGF and angiopoietin–TIE signaling, endothelial markers and hypoxia and chemokine regulators of angiogenesis (Carmeliet & Jain, 2011), and mesenchymal markers, core EMT transcription factors, matrix metalloproteinases, epithelial junction genes and TGF-β ligands and receptors for EMT (Kalluri & Weinberg, 2009; Nieto et al., 2016). The four epithelial junction genes (CDH1, OCLN, CLDN1 and TJP1) were scored in the same direction as the mesenchymal genes. Within each sample, all genes were converted to percentile ranks (ties averaged), and a program score was the mean rank of the signature genes minus the mean rank of all other genes:

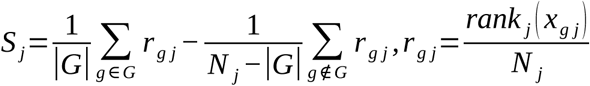

where *x_g j_* is the expression of gene g in sample j, *N _j_* is the number of genes in sample j and G is the signature gene set. The Interface score was the mean of the angiogenesis (A) and EMT (E) scores, z-standardized within each cohort, without prior standardization of the two components:

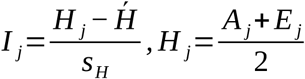

where *HỸ* and *s_H_* are the mean and standard deviation of H in the cohort. Coupling was measured as the Spearman correlation between the two program scores, and each score was compared with the mean z-scored expression of its 20 genes as an orientation check.

### Bridge-gene identification

Each of the 20,213 genes was correlated with the angiogenesis and EMT scores across the 342 tumors, with genome-wide Benjamini–Hochberg adjustment for each program. Bridge genes met all four criteria below and were ranked by the mean of their two correlations; signature genes were not excluded:

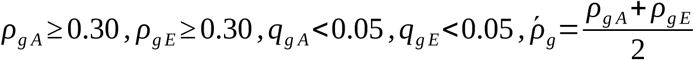

where ρ and q are the Spearman correlation and adjusted P value of gene g with the angiogenesis (A) or EMT (E) score. The number of bridge genes was tested with B = 1,000 permutations that shuffled the rows of the score table relative to the expression profiles, keeping each tumor’s two scores paired, and repeated the full selection:

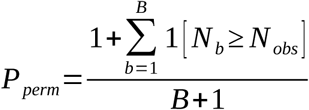

where *N _obs_* and *N_b_* are the numbers of bridge genes in the observed data and in permutation b. Stability of the 50 top-ranked bridge genes was the proportion of B = 1,000 bootstrap resamples of tumors in which both programs met the correlation and unadjusted P value criteria, without recomputing program scores:

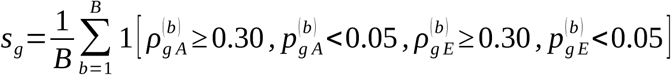

### Gene-set and pathway enrichment

Bridge genes were tested for over-representation in five predefined gene sets (one-sided Fisher’s exact tests against the 20,213 tested genes; Benjamini–Hochberg across sets): the two signatures, ECM remodeling (FN1, COL1A1, COL1A2, COL3A1, COL5A2, MMP2, MMP9, MMP14, VCAN and LAMA4), TGF-β (TGFB1, TGFB2, TGFBR1, TGFBR2, SMAD2, SMAD3, SMAD4, SNAI1 and ZEB1) and hypoxia (HIF1A, EPAS1, CA9, SLC2A1, LDHA and BNIP3). KEGG enrichment was tested separately for the bridge genes and the final candidates with the Enrichr KEGG_2021_Human library (320 pathways; Kanehisa & Goto, 2000; Kuleshov et al., 2016) retrieved with GSEApy (Fang et al., 2022). Pathways were restricted to the 20,213 tested genes, which also formed the background, and those with at least five background genes and at least one overlapping gene were tested (297 and 150 pathways, respectively) by one-sided Fisher’s exact tests with Benjamini–Hochberg adjustment (FDR < 0.05). Ten of the 107 significant bridge-gene pathways (FDR ranks 1–3, 7, 9, 11, 12, 16, 17 and 24) were selected for display. These were chosen to show non-redundant pathways from the 107 significant terms. The analysis was repeated with the GSEApy enrichr function, with the same library and background, as a cross-check.

### Tumor microenvironment adjustment

Five marker scores were calculated by converting each marker gene to a percentile rank across tumors and averaging the ranks within each tumor: CAF (COL1A1, COL1A2, COL3A1, COL5A1, COL5A2, DCN, LUM, FAP, PDGFRA, PDGFRB, THY1, ACTA2 and TAGLN), endothelial (PECAM1, VWF, EMCN, KDR, ESAM, ENG, RAMP2, PLVAP, CDH5 and EGFL7), macrophage (CD68, CSF1R, C1QA, C1QB, C1QC, CTSS, FCER1G, TYROBP, AIF1 and LST1), immune (PTPRC, CD3D, CD3E, CD247, CD79A, MS4A1, NKG7, GNLY and HLA-DRA; TRAC was not present) and hypoxia (CA9, VEGFA, SLC2A1, LDHA, PGK1, ENO1, BNIP3, P4HA1, NDRG1 and ADM). For marker set *K_k_*, the score of tumor j was

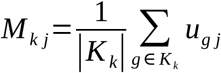

where *u_g j_* is the percentile rank of gene g in tumor j across tumors. Each marker score was correlated with the angiogenesis, EMT and Interface scores (Benjamini–Hochberg across the 15 tests). The Interface score was regressed on the five z-standardized marker scores by ordinary least squares, and the standardized residuals formed a TME-adjusted Interface score:

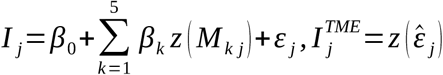

where z denotes z-standardization. A bridge gene had residual support when its correlation with the TME-adjusted Interface score reached |ρ| ≥ 0.20 with FDR < 0.05. In a program-specific analysis, the angiogenesis and EMT scores were each residualized on the marker scores in the same way, and dual residual support required ρ ≥ 0.20 with FDR < 0.05 for both; these residual correlations were compared with external-cohort correlations and provided the TME criterion for focused candidates.

### Tumor purity and partial correlations

Stromal, immune and ESTIMATE scores were calculated with the R package estimate (Yoshihara et al., 2013; filterCommonGenes with gene symbols; estimateScore with platform = “illumina”), and tumor purity was calculated from the ESTIMATE score and limited to 0–1:

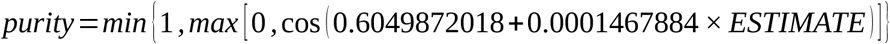

Partial Spearman correlations were computed by regressing the ranks of each variable of interest on the ranks of the k covariates *c*_1_, …, *c_k_* and correlating the residuals (Pearson), with P values from a t distribution with n − 2 − k degrees of freedom:

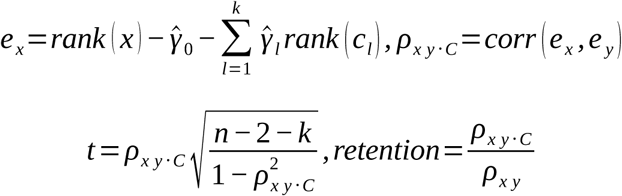

where *e _y_* is defined in the same way as *e_x_*, and retention is the ratio of the partial to the unadjusted coefficient. The angiogenesis–EMT correlation was adjusted for purity, for stromal and immune scores, and for all three. Final candidates were retained after adjustment for purity, or for purity plus stromal and immune scores, when both partial correlations were ≥ 0.20 with FDR < 0.05.

### Single-cell analysis

For each final candidate, mean expression and the percentage of cells with non-zero expression were calculated within each annotated GSE103322 compartment, with the compartment’s cell count as the denominator, and the dominant compartment was the one with the highest mean expression (ties broken by the percentage of expressing cells). An epithelial-restricted EMT score used the EMT signature genes detected in at least 10% of the 2,215 malignant cells and with non-zero variance among them (15 genes; CDH2, MMP9, TWIST1, TWIST2 and ZEB2 removed); it was recalculated in TCGA-OSCC and correlated with the angiogenesis score.

### Cell-type deconvolution

Cell-type fractions in the 342 tumors were estimated with InstaPrism, an implementation of BayesPrism (Chu et al., 2022; Hu & Chikina, 2024), with GSE103322 as the reference. Log-scale values x were converted to a linear scale for bulk data and to a TPM-like scale for single-cell data:

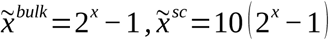

Compartments with fewer than 20 cells (myocytes) were excluded, leaving eight cell types (malignant, fibroblast, endothelial, T cell, B cell, macrophage, mast and dendritic cells), each modeled as one cell state, and the 18,523 of 19,014 shared genes with non-zero expression in both datasets were used. The reference was built with refPrepare and InstaPrism was run with filter = TRUE, pseudo.min = 1 × 10^−8^ and input_type = “refPhi_cs”; the initial posterior fractions were used and compared with ESTIMATE purity. For fraction adjustment, the angiogenesis and EMT scores were rank-residualized on all eight fractions and correlated with each final candidate’s unadjusted expression (a semi-partial correlation); candidates were retained when both correlations were ≥ 0.20 with FDR < 0.05.

### Partial-EMT score

A 15-gene partial-EMT (pEMT) signature adapted from Puram et al. (2017) was scored with the same rank-based method: SERPINE1, TGFBI, MMP10, LAMC2, P4HA2, PDPN, ITGA5, LAMA3, CDH13, TNC, MMP2, EMP3, INHBA, LAMB3 and VIM. These are the 15 common p-EMT genes listed first in the p-EMT program of Puram et al. (2017). The Interface, pEMT and purity scores were compared with pairwise and partial Spearman correlations (Interface–pEMT adjusted for purity; Interface–purity adjusted for pEMT), and the analysis was repeated with a 13-gene pEMT score excluding MMP2 and VIM, which are also EMT signature genes.

### Copy-number and methylation analyses

Copy-number and methylation data were matched to expression by patient barcode and were not adjusted for purity. GISTIC2 thresholded values (−2 to +2; Mermel et al., 2011) were available for 337 patients; duplicated gene symbols were collapsed when their rows were identical and excluded otherwise. For the 2,274 testable bridge genes, copy number was correlated with the gene’s own expression (at least 20 paired samples), and copy-number support was ρ > 0.20 with FDR < 0.05. Event frequencies were calculated for gain (≥ +1, including amplification), amplification (+2), deletion (≤ −1, including deep deletion) and deep deletion (−2). Methylation β-values outside 0–1 were set to missing, CpGs observed in at least 20% of patients (396,065 of 485,577) were kept, and remaining missing values were imputed with the CpG median. Each CpG was correlated with the Interface score, with high confidence defined as FDR < 0.05 and |ρ| ≥ 0.20, and CpGs were annotated with the UCSC_RefGene_Name field of IlluminaHumanMethylation450kanno.ilmn12.hg19; a bridge gene had methylation support when at least one annotated CpG was high-confidence, in either direction. For the final candidates, promoter CpGs (TSS1500, TSS200, 5′UTR or 1stExon) were correlated with their gene’s expression (303 CpG–gene pairs), and promoter-methylation support required ρ ≤ −0.20 with FDR < 0.05 against both the Interface score and expression.

### miRNA analyses

Mature miRNAs (miRBase MIMAT accessions) observed in at least 20% of the 314 patients with miRNA data (1,101 of 2,246) were correlated with the Interface score, with high confidence defined as FDR < 0.05 and |ρ| ≥ 0.20, and accessions were converted to mature names with miRBase release 22 (Kozomara et al., 2019). Functional human miRNA–target interactions from miRTarBase release 10.0 (Cui et al., 2025) linking high-confidence miRNAs negatively associated with the Interface score to the final candidates were tested by Spearman correlation in matched patients (3,422 pairs; at least 63 patients; samples from the same patient collapsed by their median). Pairs with ρ ≤ −0.20 and FDR < 0.05 were classed as inverse, genes with at least one inverse pair had miRNA support, and interactions supported by a reporter assay and/or Western blot were flagged as having strong experimental evidence.

### Candidate prioritization and replication

Bridge genes with both copy-number and methylation support and with TME-adjusted support were carried forward to GSE30784. There, angiogenesis and EMT scores were calculated in the 167 OSCC samples with the same scorer, and each candidate present on the array was correlated with both scores, with the relevant score recalculated without the gene for candidates that are signature genes. Final candidates had ρ ≥ 0.20 with FDR < 0.05 for both programs; a stricter subset required ρ ≥ 0.30 for both. Program scores in OSCC and normal mucosa were compared with two-sided Mann–Whitney U tests (Benjamini–Hochberg across the three scores), with rank-biserial effect sizes

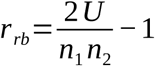

where U is the Mann–Whitney statistic for OSCC versus normal mucosa and *n*_1_ and *n*_2_ are the group sizes. Final candidates were ordered by an integrated rank score, the equal-weight mean of within-layer percentile ranks for bridge strength, copy-number–expression correlation and the strongest methylation–Interface correlation:

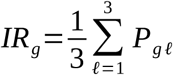

where *P_g ℓ_* is the percentile rank of gene g in layer ℓ, set to zero when the gene has no support in that layer. Of four gene-resolved evidence layers (RNA bridge, copy-number, methylation and miRNA support), only miRNA support was assessed after selection.

### Focused candidates

Ten focused candidates were drawn from the final candidates: genes mapped in all 229 GSE30784 samples, with higher median expression from normal mucosa to dysplasia and from dysplasia to OSCC, TME-adjusted support and at least one of copy-number, promoter-methylation or inverse miRNA support, were ranked by the smaller of their two median shifts, and the top 10 were kept:

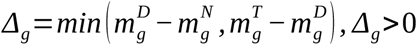

where *m^N^*, *m^D^* and *m^T^* are the medians of gene-wise z-scored expression (standardized across all 229 samples) in normal mucosa, dysplasia and OSCC. OSMR, MICAL2 and LHFPL2 were selected from these 10 as three illustrative focused candidates for detailed characterization. These were the only focused candidates with copy-number, promoter-methylation and inverse miRNA support.

### Cross-cohort and robustness analyses

Angiogenesis–EMT coupling was compared across 12 bulk expression datasets: TCGA-OSCC, GSE30784 (all 229 samples), GSE37991 (n = 80), GSE41613 (n = 97) and eight TCGA cancer types (BRCA, COAD, GBM, LUAD, LUSC, PAAD, READ and STAD). Program scores were calculated in each dataset using all samples in its expression matrix, including normal samples (4 to 114 per TCGA cancer type; normal mucosa and dysplasia in GSE30784); these values were computed with an earlier version of the scoring pipeline and are not reproduced exactly by the scorer described above, and P values were Benjamini–Hochberg adjusted across the 12 datasets. Coupling was also recalculated with the MSigDB Hallmark angiogenesis and epithelial–mesenchymal transition sets (v2023.1; Liberzon et al., 2015; 36 and 194 genes present) replacing either or both signatures, after leaving out each signature gene in turn, and for 1,000 random signature pairs that matched each signature’s gene count within 20 mean-expression bins, excluded the 40 signature genes and did not overlap within a pair; the empirical P value was (k + 1)/(N + 1), where k is the number of the N = 1,000 pairs with coupling at least as strong as observed. Final candidates persisted when both correlations with the Hallmark-based scores were ≥ 0.20 with FDR < 0.05.

### CPTAC-HNSCC analysis

In the 110 CPTAC-HNSCC tumors, program scores were calculated from RNA-seq with the same scorer, and measured final candidates replicated when both program correlations were ≥ 0.20 with FDR < 0.05. For OSMR, MICAL2 and LHFPL2, protein abundance was correlated with the RNA-based Interface score and with the gene’s own mRNA in tumors with non-missing protein values (at least 8), without multiple-testing correction. CPTAC-HNSCC was analyzed as a mixed-site HNSCC comparison cohort rather than as an independent OSCC validation cohort.

### Clinical and survival analyses

Pathologic stage was grouped as I–IV (substages merged) and as early (I–II) versus advanced (III–IV), and nodal status as N0 versus N+ (N1–N3), excluding NX and missing values. Interface scores were compared across stages (Kruskal–Wallis), between N0 and N+ tumors (two-sided Mann–Whitney U), against ordinal stage (Spearman) and between early and advanced stage, with Benjamini–Hochberg adjustment across the four tests. Overall survival was defined by death from any cause, with time to death or last follow-up; non-positive times were excluded, leaving 340 patients with 154 deaths. Kaplan–Meier curves split at the median Interface score were compared with the log-rank test; the continuous Cox model was the primary analysis. Cox models were fitted with statsmodels PHReg (Seabold & Perktold, 2010) using Efron’s method for ties, for the Interface score alone and adjusted for age, sex and advanced stage or for age, sex and N+ status, with the Interface score and age z-standardized within each complete-case model. The stage-adjusted model was

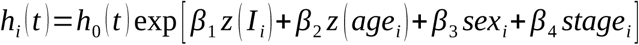

where *stage_i_* indicates advanced stage (replaced by N+ status in the nodal-adjusted model) and exp (*β*_1_) is the hazard ratio per 1-SD increase in the Interface score. Proportional hazards in the adjusted models were assessed with Schoenfeld-residual tests in lifelines (Davidson-Pilon, 2019), with age per 10 years, Kaplan–Meier time transformation and Benjamini–Hochberg adjustment across covariates.

### Statistical analysis and software

Unless stated otherwise, associations were measured with Spearman’s rank correlation (average ranks for ties), with two-sided P values from the t approximation with n − 2 degrees of freedom, computed with SciPy (Virtanen et al., 2020) or an equivalent vectorized implementation:

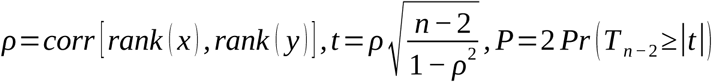

Over-representation tests were one-sided and all other tests two-sided. Multiple testing was controlled with the Benjamini–Hochberg procedure (Benjamini & Hochberg, 1995) within each family of m tests, with adjusted P values computed from the ordered P values *p*_(1)_ ≤ … ≤ *p*_(_ *_m_*_)_ as

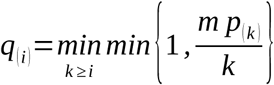

and FDR < 0.05 considered significant; thresholds and correction families are listed in Supplementary Table S1. Bootstrap and random-signature sampling used NumPy default_rng with seed 42, permutations used one generator per permutation from NumPy SeedSequence([42, permutation index]), and deconvolution used set.seed(42) in R. Gene symbols were used as provided by each dataset. Analyses were performed in Python 3.12.5 with NumPy 2.2.3–2.2.6, pandas 2.2.3, SciPy 1.15.2, statsmodels 0.14.6, scikit-learn 1.8.0, lifelines 0.30.3 and GSEApy 1.1.13, and in R with the estimate and InstaPrism packages. The R and package versions used for these steps were not recorded.

## Results

### A coordinated angiogenesis–EMT transcriptional state delineates a bridge-gene framework in OSCC

To determine the coupling between angiogenesis and EMT, gene-expression profiles from 342 primary oral squamous cell carcinoma (OSCC) tumors in the oral cavity subset of TCGA-HNSC (hereafter, TCGA-OSCC) were analyzed. The analysis used two predefined, non-overlapping custom signatures of 20 genes each. Program scores were calculated using a rank-based sample-scoring approach. A strong correlation was observed between the angiogenesis and EMT programs (Spearman ρ = 0.809, P = 2.7 × 10^−80^; Fig. 1A). This demonstrates that tumors with higher angiogenesis program scores also exhibit higher EMT program scores.

**Figure 1.**
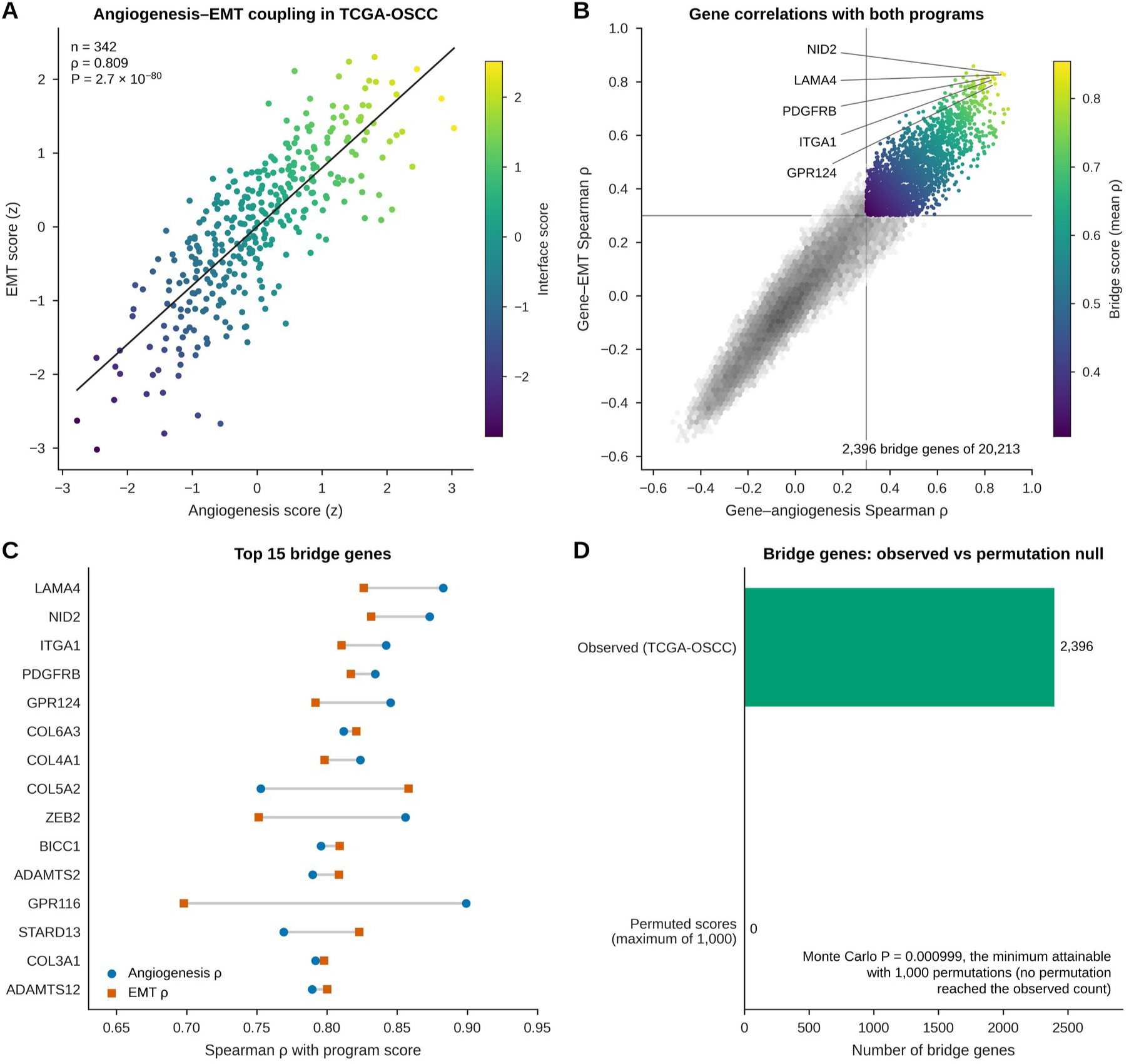
Angiogenesis–EMT coupling and bridge genes in TCGA-OSCC. (A) Angiogenesis and EMT program scores (z-standardized) in 342 tumors, colored by Interface score; the line is a least-squares fit. (B) Spearman correlation of each of 20,213 genes with the angiogenesis and EMT scores. Gray lines mark ρ = 0.30; colored points are the 2,396 bridge genes (ρ ≥ 0.30 and FDR < 0.05 for both programs), colored by bridge score (mean ρ). (C) Correlations of the 15 top-ranked bridge genes with each program score. (D) Number of bridge genes observed and the maximum number obtained in 1,000 permutations of the score table.

To further determine whether the angiogenesis–EMT coupling extends beyond the predefined signature gene level, we performed a transcriptome-wide correlation analysis, in which the expression level of every gene was correlated with angiogenesis and EMT scores across the same tumor samples. Genes showing positive correlations with both programs were defined as bridge genes. We applied predefined criteria to classify a gene as a bridge gene when its expression was positively correlated with each program score (Spearman ρ ≥ 0.30 for each) and both associations remained significant after Benjamini–Hochberg multiple-testing correction (FDR < 0.05). We identified 2,396 genes that met these criteria and classified them as bridge genes, including 31 of the 40 signature genes (17 of 20 angiogenesis genes and 14 of 20 EMT genes). These results show that genes associated with both programs extend beyond the predefined signatures (Fig. 1B).

We next sought to identify the top bridge-gene transcripts that show the strongest positive coupling with the angiogenesis and EMT program scores. Bridge genes were ranked using a mean Spearman correlation measure of their associations with the two programs. Among the strongest bridge-gene transcripts, LAMA4, NID2, ITGA1, PDGFRB, and ADGRA2 (GPR124) showed strong positive correlations with both programs (Fig. 1C).

After determining the bridge genes from the scores, we evaluated whether the bridge-gene architecture was statistically non-random. To this end, we compared the bridge-gene pipeline against a permuted null distribution in which program scores were randomly permuted against sample expression profiles across 1,000 Monte Carlo iterations, using the same dual-correlation thresholds (ρ ≥ 0.30 and FDR < 0.05 for both programs). Fig. 1D shows that no bridge genes were identified in any of the 1,000 iterations (Monte Carlo P = 0.000999, the minimum attainable P). This result supports a non-random pattern of shared expression under the permutation scheme used: the observed number of genes associated with both program scores was not reproduced after random reassignment of the scores to the expression profiles.

To further characterize the underlying architecture of these transcriptional states, we verified score orientation (Fig. 2A), examined score distributions (Fig. 2B), tested pathway over-representation (Fig. 2C), and assessed resampling stability (Fig. 2D) in the TCGA-OSCC cohort. First, as a signature-orientation check, we correlated each program score with the mean standardized expression of its 20 signature genes from RNA-seq data, using Spearman rank correlation. Fig. 2A shows positive correlations, with ρ = 0.983 for the angiogenesis score and ρ = 0.977 for the EMT score, confirming that each score is oriented in the same direction as the expression of its signature genes.

**Figure 2.**
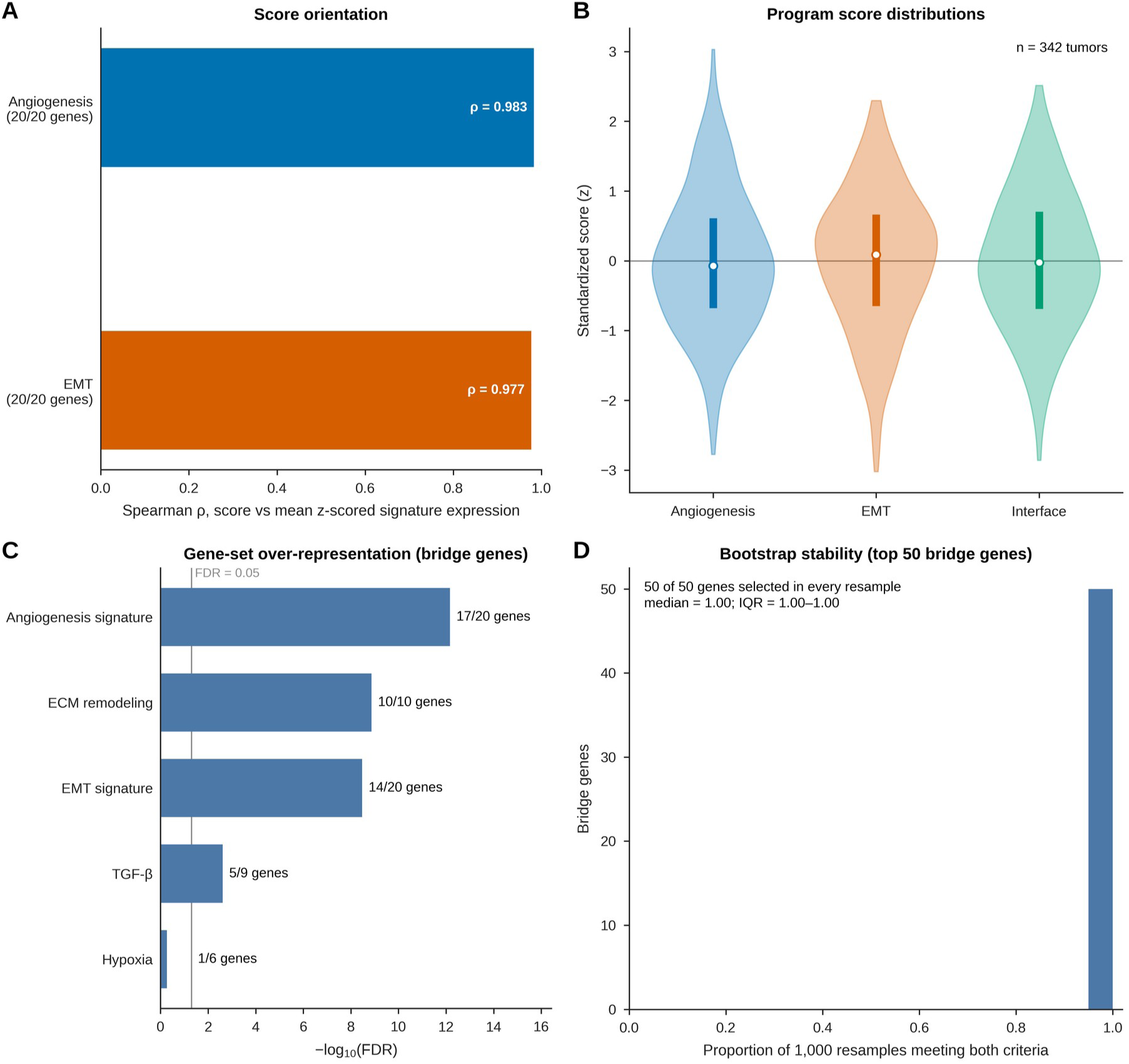
Score orientation, score distributions, gene-set over-representation and bootstrap stability. (A) Spearman correlation of each program score with the mean z-scored expression of its 20 signature genes. (B) Distributions of the standardized angiogenesis, EMT and Interface scores (n = 342); points show medians and bars interquartile ranges. (C) Over-representation of five predefined gene sets among the bridge genes (one-sided Fisher’s exact tests, Benjamini–Hochberg adjusted); labels give the number of set genes among the bridge genes, and the gray line marks FDR = 0.05. (D) Proportion of 1,000 bootstrap resamples in which each of the top 50 bridge genes met both criteria.

To further test how these transcriptional states vary across individual primary OSCC tumors, we evaluated the angiogenesis, EMT, and angiogenesis–EMT Interface program scores and their density distributions (Fig. 2B). The Interface score was defined as the z-standardized mean of the angiogenesis and EMT program scores. These standardized scores showed broad, unimodal distributions across individual tumors, which supports inter-tumor heterogeneity and describes angiogenesis, EMT, and Interface as continuous quantitative transcriptional states.

Given this inter-tumor variation, we next investigated which biological processes are represented by this bridge-gene architecture. We therefore tested the 2,396 bridge genes for over-representation of a predefined panel of biological gene sets. The bridge genes were significantly enriched for the angiogenesis, extracellular matrix remodeling, EMT, and TGF-β gene sets, but not for the hypoxia gene set (FDR = 0.531; Fig. 2C). Because the bridge genes were selected for their correlation with the angiogenesis and EMT scores, enrichment of those two gene sets is expected by construction.

Finally, to test whether the dual association status of the top-ranked bridge genes is stable under resampling of tumors, we performed 1,000 bootstrap resamples. We evaluated the selection frequency and dual-correlation stability for the top 50 bridge genes, whose median bootstrap stability was 1.00 (IQR, 1.00–1.00; Fig. 2D). Their high selection frequency and stable dual correlations support the resampling stability of these 50 genes.

### Angiogenesis–EMT coupling is attenuated but retained after accounting for tumor composition and cellular context

Because bulk transcriptomic data reflect both malignant epithelial cells and non-malignant elements, we sought to assess whether the angiogenesis–EMT coupling remains detectable after accounting for tumor purity and non-malignant cell-type scores. To evaluate this, we specifically determined which non-malignant tumor microenvironment (TME) cell types show the strongest correlation with the angiogenesis, EMT, and Interface program scores. The results in Fig. 3A show strong correlations between cancer-associated fibroblast (CAF) marker scores and angiogenesis (ρ = 0.80), EMT (ρ = 0.81), and Interface (ρ = 0.84). Additionally, endothelial scores were strongly associated with angiogenesis (ρ = 0.82) and Interface (ρ = 0.73). Moderate correlations were observed for macrophage and immune signatures, whereas hypoxia showed minimal association. These results identify fibroblast and endothelial marker scores as prominent correlates of the program scores. After observing the associations with CAF and endothelial marker scores, we evaluated whether the bridge-gene signal remained detectable after adjustment for TME-related variables. Comparing each bridge gene’s original bridge score with its correlation with the TME-adjusted Interface score identified a subset with retained positive associations and a smaller subset with negative associations after adjustment (Fig. 3B).

**Figure 3.**
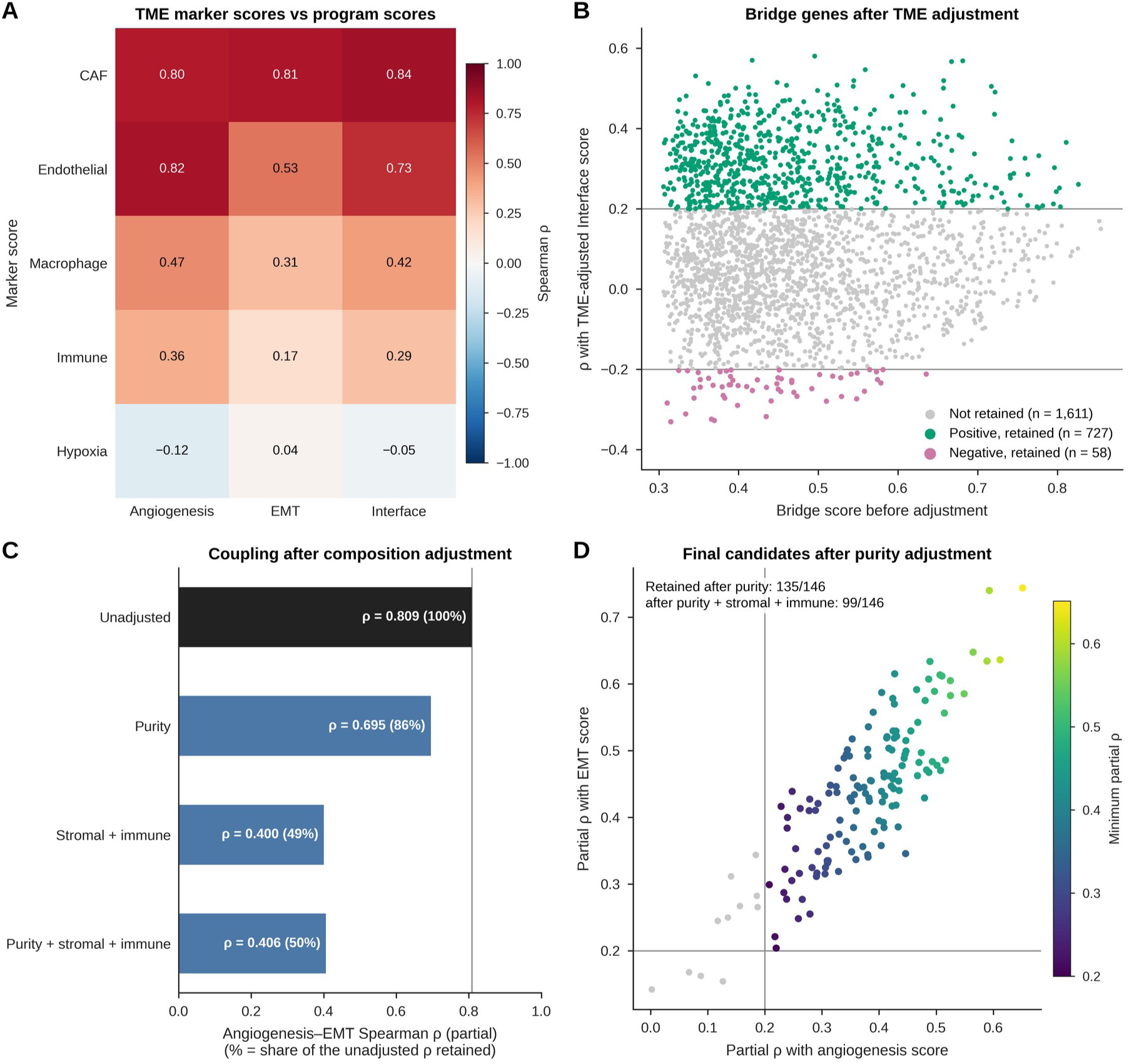
Tumor microenvironment marker scores and composition adjustment. (A) Spearman correlations of five marker scores with the angiogenesis, EMT and Interface scores. (B) Correlation of each bridge gene with the TME-adjusted Interface score against its bridge score before adjustment; lines mark ρ = ±0.20, and colored points are retained positive and negative associations (|ρ| ≥ 0.20, FDR < 0.05). (C) Angiogenesis–EMT Spearman ρ without adjustment and partial ρ after adjustment for purity, for stromal and immune scores, and for all three; percentages give the share of the unadjusted ρ retained. (D) Partial correlations of the 146 final candidates with each program score after purity adjustment, colored by the smaller partial ρ; gray points were not retained.

Next, we evaluated how the correlation between the program scores changed after adjustment for estimated tumor purity and stromal and immune infiltration. The unadjusted angiogenesis– EMT correlation was ρ = 0.809. After purity adjustment, the correlation decreased to ρ = 0.695, retaining approximately 86% of its original magnitude. Adjustment for stromal and immune infiltration reduced the correlation to ρ = 0.400, corresponding to 49% retention. Following joint adjustment, the correlation remained positive at ρ = 0.406, retaining approximately 50% of the unadjusted correlation magnitude (Fig. 3C). These results show attenuation of the association after adjustment for the included composition variables.

After purity adjustment, we evaluated which of the 146 final candidates (prioritized bridge genes that were also supported in the external GSE30784 cohort, as described below) still maintained a dual positive partial correlation with both program scores. Fig. 3D shows that a substantial subset of the final candidates retained positive associations with both programs after purity adjustment in TCGA-OSCC samples.

We further analyzed single-cell RNA-seq data from oral cavity tumors (Puram et al., 2017; GSE103322) to evaluate the compartment-specific expression patterns of the top bridge genes and to examine coupling using an epithelial-restricted EMT score and after cell-fraction adjustment. First, we determined which single-cell compartments are most represented by the bridge genes. Fig. 4A shows that, among the 146 final candidates, the most frequent dominant compartment was endothelial cells (n = 41), followed by fibroblasts (n = 26), malignant epithelial cells (n = 20), and macrophages (n = 18), indicating compartmental heterogeneity.

**Figure 4.**
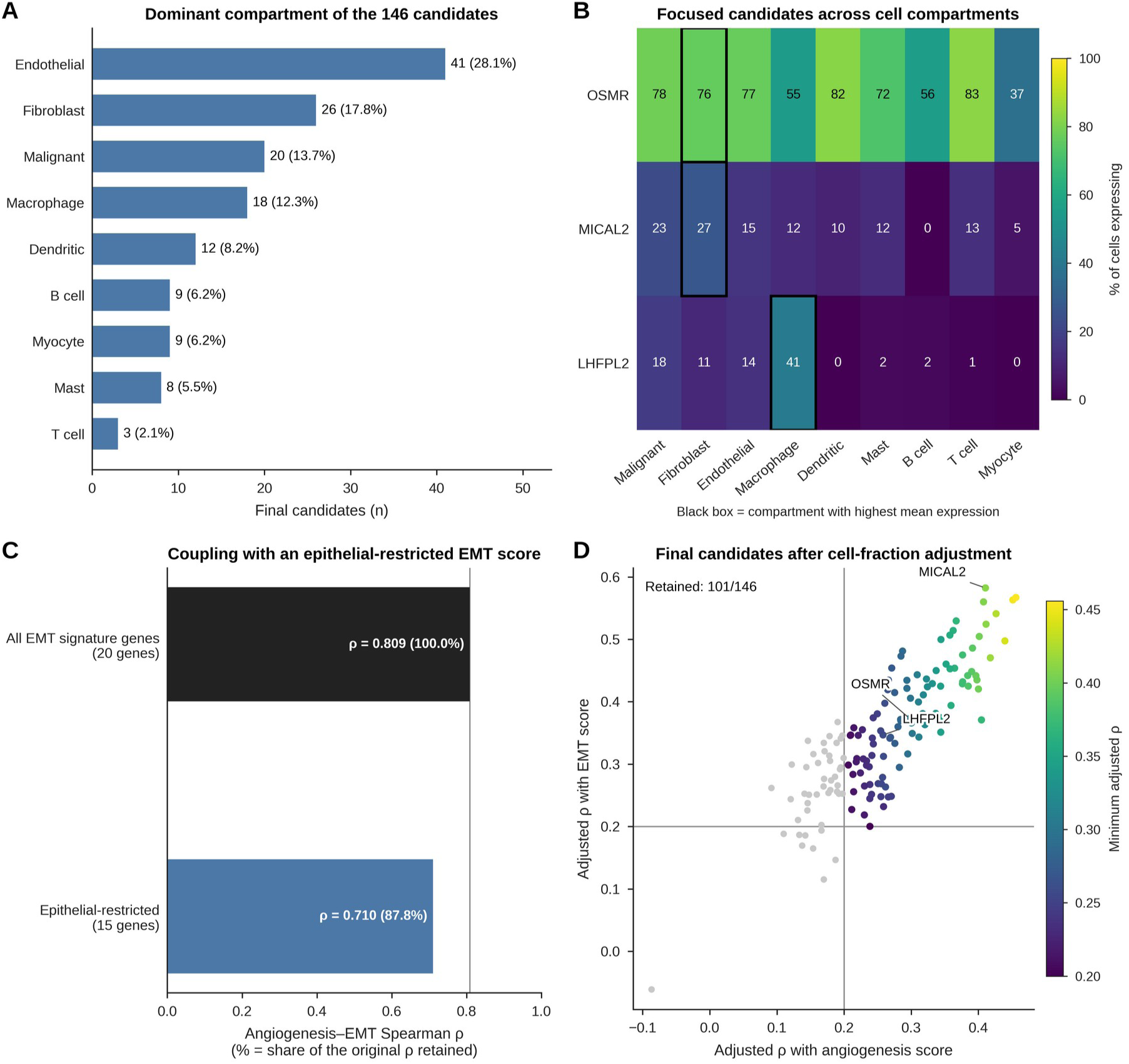
Single-cell context of the final candidates and adjustment for inferred cell fractions. (A) Dominant GSE103322 compartment (highest mean expression) of the 146 final candidates. (B) Percentage of cells expressing OSMR, MICAL2 and LHFPL2 in each compartment; black boxes mark the compartment with the highest mean expression. (C) Angiogenesis–EMT coupling with the 20-gene EMT score and with the epithelial-restricted 15-gene EMT score; percentages give the share of the original ρ retained. (D) Correlations of the final candidates with the angiogenesis and EMT scores after rank-residualization of the scores on the eight InstaPrism cell fractions, colored by the smaller adjusted ρ; gray points were not retained.

After mapping the broader compartmental distribution across the bridge-gene set, we evaluated the single-cell expression profiles of OSMR, MICAL2, and LHFPL2, three illustrative focused candidates selected for detailed characterization. Fig. 4B shows expression of the focused candidates across different cellular compartments. Fibroblasts were the dominant compartment for OSMR and MICAL2, with 76.3% and 27.0% of fibroblasts expressing each gene, respectively, whereas macrophages were the dominant compartment for LHFPL2 (40.8% of macrophages expressing). These distributions place candidate expression within stromal and myeloid compartments without indicating exclusive expression in those compartments.

Given the expression of several candidates in non-malignant compartments, we examined coupling using the epithelial-restricted EMT score. In this analysis, the correlation decreased from ρ = 0.809 to ρ = 0.710, retaining 87.8% of the baseline correlation magnitude (Fig. 4C).

To complement this epithelial-restricted program analysis, we then analyzed whether individual final candidates still maintained dual-program support after adjustment for single-cell-derived inferred cell fractions. Fig. 4D shows that 101 of 146 final candidates (69.2%) retained dual-program support after this adjustment. Additionally, the focused candidates OSMR, MICAL2, and LHFPL2 maintained a dual-program association. These results identify candidates with retained dual-program associations after adjustment for the inferred cell fractions.

After mapping the single-cell compartment distribution of the candidates, we examined deconvolution-estimated cell fractions, ESTIMATE tumor purity, and partial EMT (pEMT) scores based on a 15-gene signature (adapted from Puram et al., 2017). ESTIMATE provides an expression-based estimate of tumor purity. Fig. 5A shows that the deconvolution-estimated malignant cell fraction correlated positively with ESTIMATE tumor purity (ρ = 0.70), whereas all estimated non-malignant cell fractions correlated negatively, most strongly those of macrophages (ρ = −0.87), fibroblasts (ρ = −0.58), and T cells (ρ = −0.52). These associations indicate agreement between the inferred cell fractions and the expression-based purity estimate.

**Figure 5.**
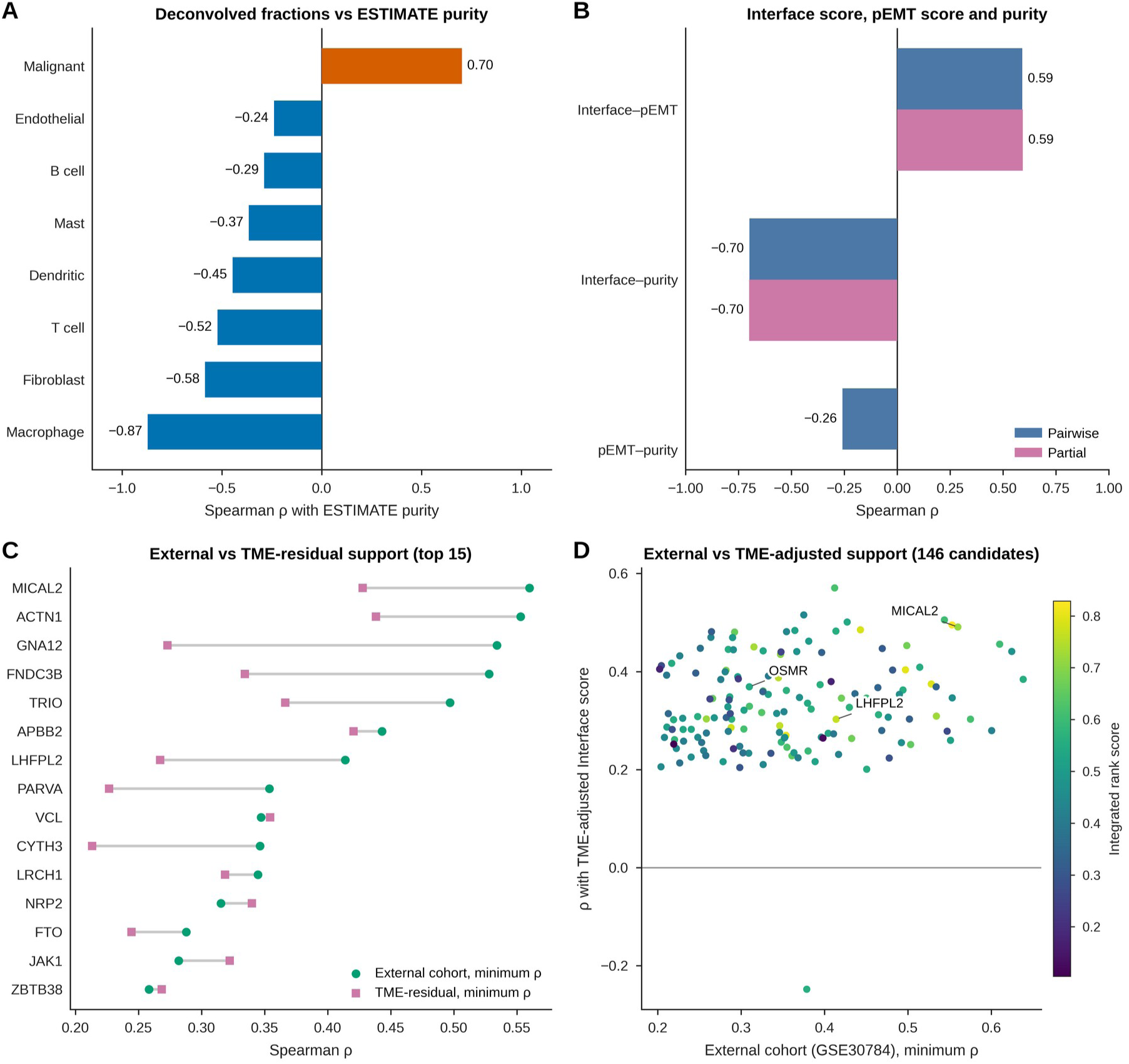
Cell fractions, partial-EMT score and TME-residual support. (A) Spearman correlation of each InstaPrism-estimated cell fraction with ESTIMATE purity. (B) Pairwise and partial Spearman correlations among the Interface score, the 15-gene pEMT score and purity (Interface–pEMT adjusted for purity; Interface–purity adjusted for pEMT). (C) Minimum ρ in GSE30784 (external cohort) and minimum TME-residual ρ for 15 final candidates. (D) Minimum GSE30784 ρ against the correlation with the TME-adjusted Interface score for the 146 final candidates, colored by integrated rank score.

The Interface score correlated positively with the pEMT score (ρ = 0.59) and negatively with estimated tumor purity (ρ = −0.70), whereas pEMT and purity were only weakly correlated (ρ = −0.26). Both associations were unchanged in partial correlation analyses (Interface–pEMT adjusted for purity, partial ρ = 0.59; Interface–purity adjusted for pEMT, partial ρ = −0.70; Fig. 5B). The pEMT signature shares two genes with the custom EMT signature (MMP2 and VIM). After these genes were removed from the pEMT score, the Interface–pEMT association persisted (ρ = 0.554; purity-adjusted partial ρ = 0.567), indicating that the observed relationship was not explained solely by direct gene-set overlap.

Further, for 15 final candidates, we compared support in the external cohort (GSE30784; described below) with support after TME adjustment, plotting the external-cohort minimum ρ against the TME-residual minimum ρ (Fig. 5C). These comparisons distinguish support from external-cohort associations from support after adjustment for the included TME variables.

Next, we compared the external-cohort minimum ρ with the TME-residualized Interface ρ across the final candidates, which showed that a subset of genes exhibits strong correlations on both axes (Fig. 5D). These analyses identify candidates supported by external-cohort associations and by associations retained after adjustment for the included TME variables.

### Copy-number, methylation, and miRNA analyses provide complementary candidate-level support

We next examined whether bridge-gene expression was associated with copy-number variation (CNV) and promoter methylation patterns. To assess this, we evaluated how RNA expression was associated with CNV dosage in primary OSCC tumors. Fig. 6A indicates that, among 2,274 tested bridge genes, 553 exhibited statistically significant positive CNV–RNA associations, including ADD1 (ρ = 0.70), SRPR (ρ = 0.69), ARCN1 (ρ = 0.68), PICALM (ρ = 0.68), and SNX19 (ρ = 0.65), supporting dosage-associated expression for a subset of the tested bridge genes. Because this analysis relates each gene’s copy number to its own expression, it provides candidate-level genomic support rather than a direct test of the Interface.

**Figure 6.**
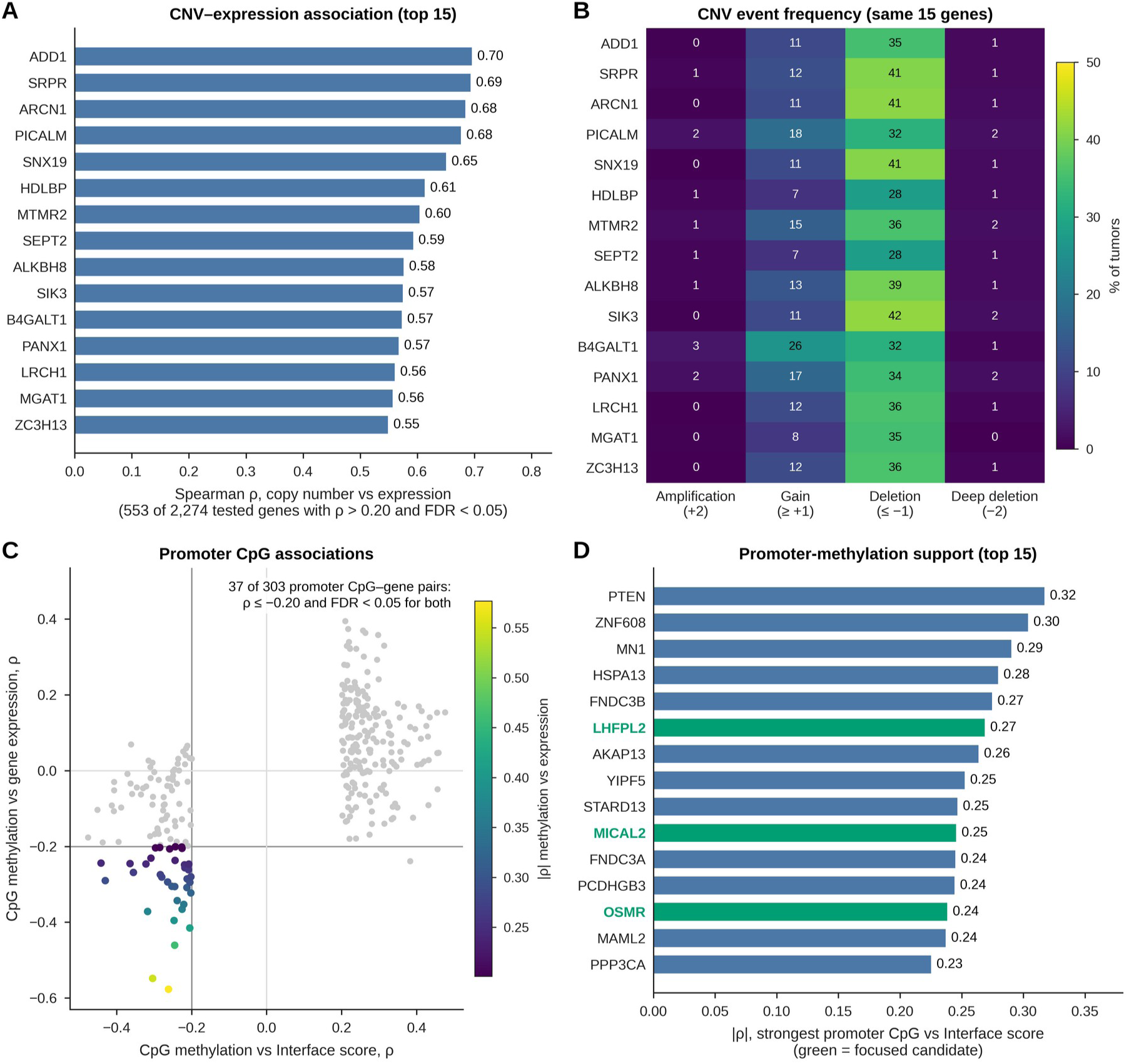
Copy-number and promoter-methylation support. (A) The 15 bridge genes with the strongest Spearman correlations between copy number and expression. (B) Percentage of tumors with amplification, gain, deletion and deep deletion for the same genes. (C) Correlation of promoter CpG methylation with the Interface score against its correlation with the gene’s expression for the final candidates; colored points are pairs with ρ ≤ −0.20 and FDR < 0.05 for both. (D) The 15 candidates with the strongest promoter CpG–Interface association; focused candidates are shown in green.

In addition, we examined the spectrum of CNV event types, specifically shallow deletions, deep deletions, gains, and amplifications, among bridge genes with positive CNV–RNA associations. Fig. 6B shows that shallow deletions were more frequent than deep deletions or amplifications, whereas gains occurred at moderate-to-low frequency.

We next analyzed promoter DNA methylation by identifying CpG loci with inverse associations between methylation and both the corresponding gene’s RNA expression and the Interface score. At these loci, lower promoter methylation (hypomethylation) was associated with higher gene expression and higher Interface scores (Fig. 6C).

Next, we identified which candidate bridge genes exhibit the strongest promoter methylation support. Fig. 6D shows that the top-ranked genes PTEN, ZNF608, MN1, and HSPA13, as well as the focused candidates LHFPL2, MICAL2, and OSMR, have promoter methylation associations alongside transcriptomic support.

We further extended our association analysis to epigenome-wide CpG methylation patterns and microRNA (miRNA) expression levels, assessing their associations with the angiogenesis–EMT Interface score across OSCC tumors. Of 396,065 CpGs tested, 80,197 showed high-confidence associations with the Interface score, including both positive and negative correlations, with positive correlations more frequent (Fig. 7A).

**Figure 7.**
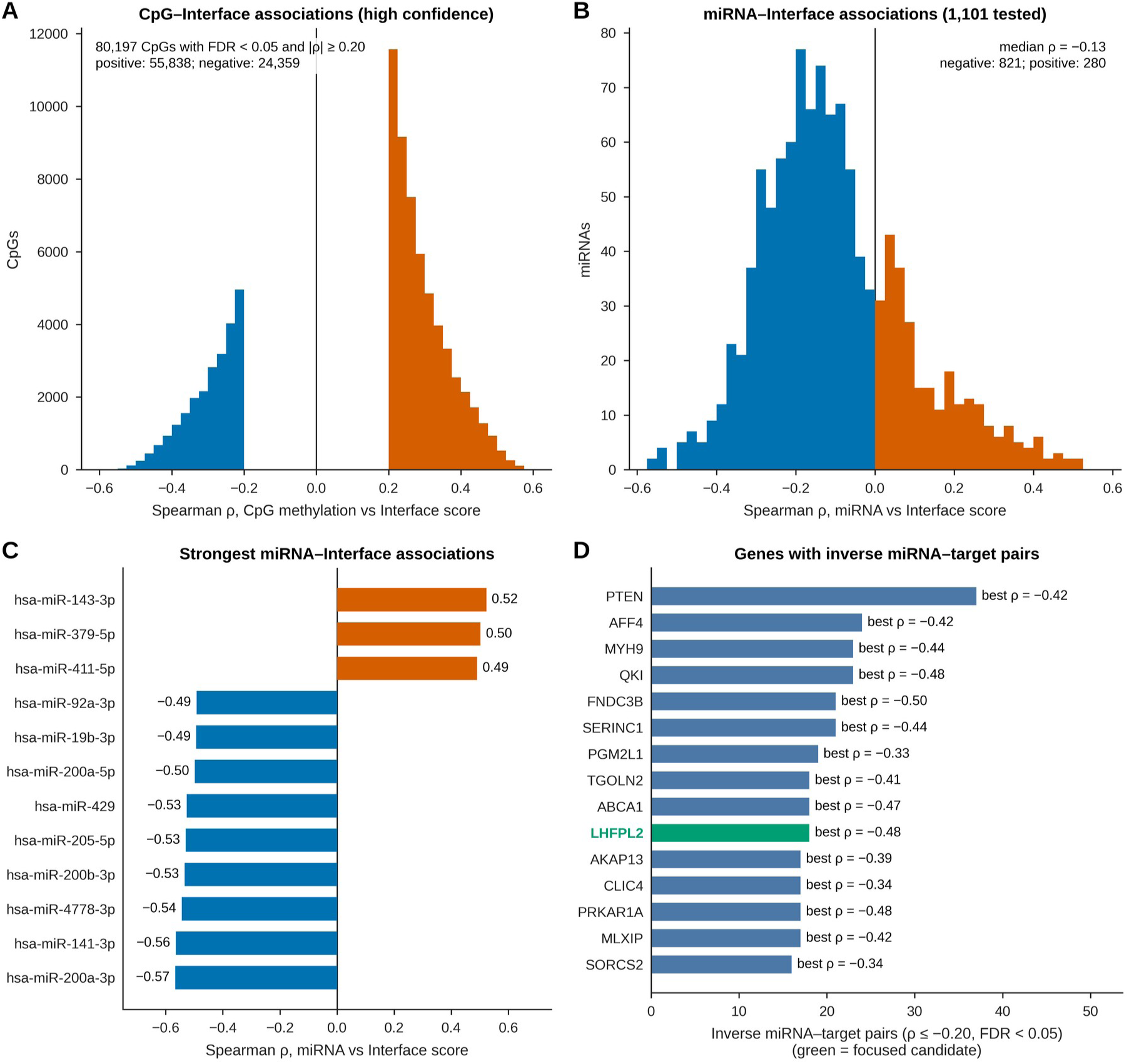
DNA methylation and miRNA associations with the Interface score. (A) Distribution of Spearman ρ for the 80,197 high-confidence CpGs (FDR < 0.05 and |ρ| ≥ 0.20). (B) Distribution of miRNA–Interface ρ for the 1,101 tested miRNAs. (C) miRNAs with the strongest positive and negative associations with the Interface score. (D) Genes with the largest numbers of inverse miRNA–target pairs (ρ ≤ −0.20, FDR < 0.05); labels give the strongest pair, and the focused candidate is shown in green.

Additionally, we examined the distribution of correlations between miRNA expression and Interface scores across OSCC samples. Across the 1,101 evaluated miRNAs, the distribution of miRNA–Interface correlation coefficients was shifted toward negative values (Fig. 7B), indicating that many miRNAs showed lower expression in tumors with higher Interface scores. Of the 1,101 miRNAs, 537 were significantly associated with the Interface score (FDR < 0.05), and 401 met high-confidence criteria.

Moreover, we examined the miRNAs that correlated negatively with the Interface score. Fig. 7C shows key epithelial miRNAs, including hsa-miR-200a-3p, hsa-miR-141-3p, hsa-miR-200b-3p, hsa-miR-429, and hsa-miR-205-5p, which also show negative correlations. A smaller subset of miRNAs exhibited positive correlations with the Interface score. Because the Interface score is strongly and inversely associated with tumor purity (Fig. 5B), these bulk miRNA associations, like the methylation associations above, may partly reflect cellular composition.

Furthermore, we evaluated the candidate bridge genes that display the strongest negative associations with predicted targeting miRNAs. Of 3,422 candidate miRNA–gene pairs tested, 916 showed qualifying inverse associations. Fig. 7D shows that PTEN had the largest number of inverse miRNA–target pairs at FDR < 0.05 (a count that grows with the number of miRNAs predicted to target a gene), whereas the strongest individual inverse correlations included those of FNDC3B (ρ = −0.50), QKI (ρ = −0.48), and LHFPL2 (ρ = −0.48). These associations identify predicted miRNA–gene pairs in which lower miRNA expression coincides with higher expression of the candidate gene.

In addition, we integrated the epigenomic, genomic, TME, and external-cohort evidence with predicted miRNA-targeting networks to prioritize genes associated with the angiogenesis–EMT Interface. Fig. 8A shows miRNA–target pairs with inverse expression correlations; among pairs with strong experimental evidence in miRTarBase (covering 15 genes), the strongest included hsa-miR-200b-3p–ETS1, hsa-miR-17-5p–JAK1, hsa-miR-17-5p–ABCA1, hsa-miR-19a-3p–ABCA1, and hsa-miR-93-5p–LATS2. Together with the observed expression associations, these links identify candidate post-transcriptional regulatory relationships for further investigation.

**Figure 8.**
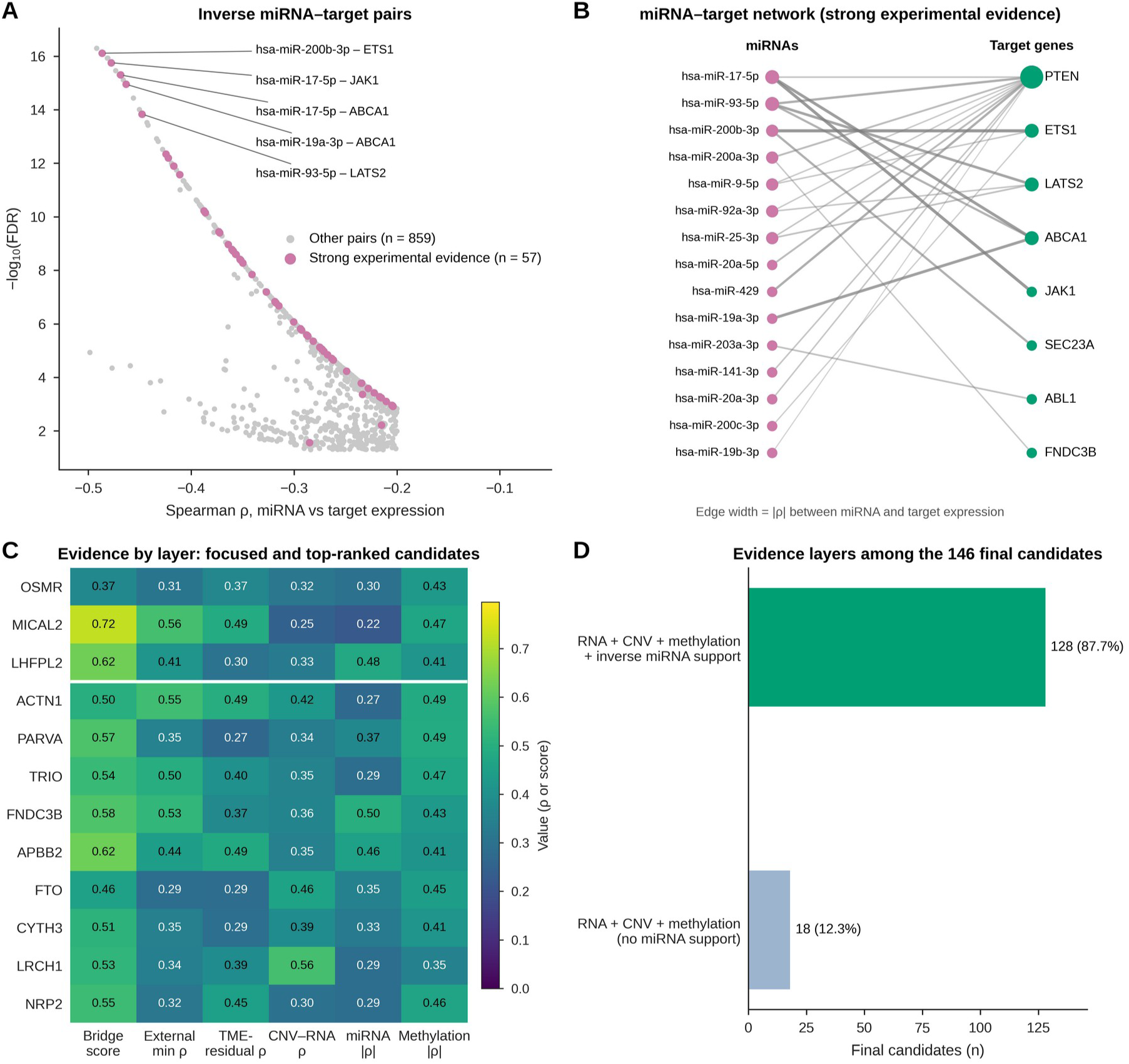
miRNA–target pairs and evidence layers of the final candidates. (A) Spearman ρ and −log10(FDR) of the 916 inverse miRNA–target pairs; pink points are pairs with strong experimental evidence in miRTarBase (reporter assay and/or Western blot), and the five strongest are labeled. (B) Network of the miRNA–target pairs with strong experimental evidence; edge width is |ρ|. (C) Evidence-layer values for the focused candidates and top-ranked final candidates. (D) Combinations of evidence layers among the 146 final candidates.

Moreover, we evaluated how candidate miRNAs and bridge genes are organized within a network of predicted miRNA–target relationships. Fig. 8B presents an interconnected, complex architecture in which multiple miRNAs connect to several candidate bridge genes (PTEN, ABCA1, ABL1, ETS1, LATS2, FNDC3B, JAK1, and SEC23A), illustrating the topology of the predicted miRNA–target network.

Next, we assessed evidence-layer support among the final candidates. Within the final candidate set, 128 of 146 genes also showed gene-resolved inverse-miRNA support in addition to the RNA, CNV, and methylation criteria used in candidate prioritization; the remaining 18 met these three criteria without qualifying miRNA support (Fig. 8C,D). The final candidate set includes the focused candidates OSMR, MICAL2, and LHFPL2.

### External-cohort replication of the angiogenesis–EMT transcriptional Interface state

Next, we examined whether angiogenesis–EMT coupling was reproduced in an external OSCC cohort. The positive angiogenesis–EMT association was reproduced in the 167 OSCC samples of GSE30784 (ρ = 0.648; Fig. 9A).

**Figure 9.**
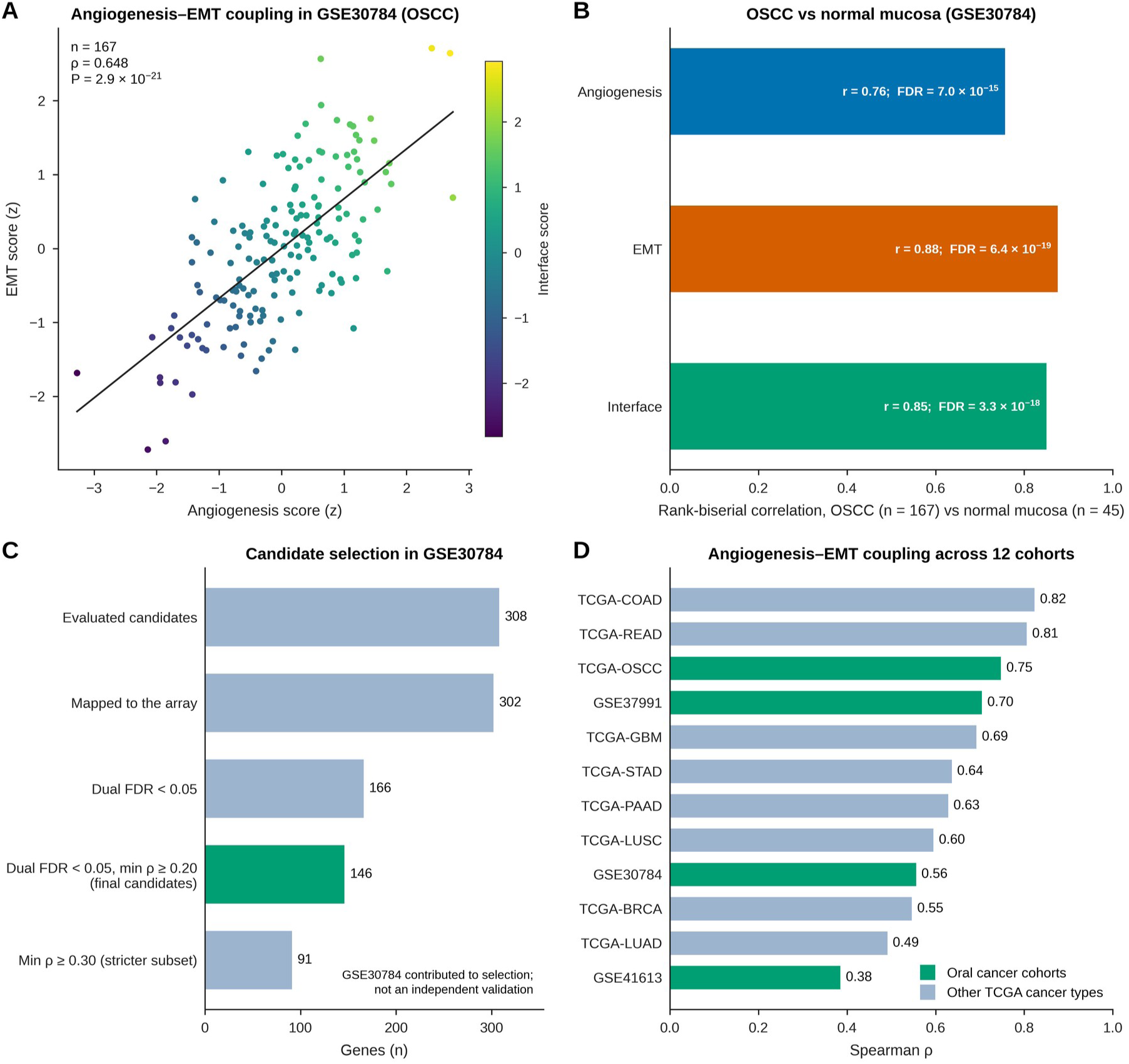
External-cohort and cross-cancer analyses. (A) Angiogenesis and EMT scores (z-standardized) in the 167 OSCC samples of GSE30784, colored by Interface score. (B) Rank-biserial correlations comparing OSCC (n = 167) with normal oral mucosa (n = 45) (two-sided Mann–Whitney U tests, Benjamini–Hochberg adjusted). (C) Selection of the final candidates in GSE30784; GSE30784 contributed to candidate selection and is not an independent validation of the candidates. (D) Angiogenesis–EMT Spearman ρ in 12 bulk expression datasets. Panel D values were computed with an earlier version of the scoring pipeline (see Methods).

Moreover, we evaluated whether program-level scores are higher in primary OSCC tumors than in non-malignant oral mucosal tissue. Fig. 9B shows that angiogenesis, EMT, and Interface scores were higher in OSCC, with large rank-biserial effect sizes of 0.76 for angiogenesis (FDR = 7.0 × 10^−15^), 0.88 for EMT (FDR = 6.4 × 10^−19^), and 0.85 for Interface (FDR = 3.3 × 10^−18^). These results show higher program scores in the analyzed OSCC tissues than in non-malignant oral mucosal tissues.

In addition to the program-level association, we examined candidate-gene associations in the external dataset. The 308 evaluated genes were bridge genes with combined RNA, CNV, and methylation support (467 genes) that also showed a residual association with the Interface after TME-marker adjustment (|residual ρ| ≥ 0.20, FDR < 0.05; two of these residual associations were negative). Of these 308 genes, 302 mapped to the external dataset. Of these, 166 met the dual FDR < 0.05 criterion, 146 met the dual FDR < 0.05 criterion with a minimum correlation of ρ ≥ 0.20, and 91 met the stricter threshold of ρ ≥ 0.30 (Fig. 9C). These results identify a subset of evaluated genes associated with both programs in the external dataset. These 146 genes form the final candidate set analyzed throughout this study. Because GSE30784 contributed to their selection, this candidate-level step is part of candidate selection rather than independent validation.

Finally, we evaluated angiogenesis–EMT coupling across 12 cancer cohorts, including eight additional TCGA cancer types. Positive associations were observed in all 12 cohorts, with ρ ranging from 0.38 to 0.82 (Fig. 9D).

### Supporting transcriptomic and proteomic evidence for prioritized candidate genes

After examining transcriptomic-level program coupling across different cohorts, we assessed protein-level support using matched RNA and mass-spectrometry proteomic profiles from the CPTAC head and neck squamous cell carcinoma (CPTAC-HNSCC) cohort, which aids in integrating proteomic evidence into the final multi-omic prioritization framework. The analysis was restricted to tumor profiles (n = 110; oral cavity, 49; larynx, 49; oropharynx, 6; lip, 4; hypopharynx, 2). Fig. 10A shows each measurable final candidate’s RNA correlations with the CPTAC angiogenesis and EMT scores; 60 of 142 measurable candidates met the dual RNA replication criterion. At the program level, RNA-based angiogenesis and EMT scores were positively correlated (ρ = 0.655, n = 110, P = 8.1 × 10^−15^; Fig. 10B). Fig. 10B also shows protein–Interface correlations and mRNA–protein concordance for the focused candidates MICAL2 (ρ = 0.515 protein–Interface; ρ = 0.726 mRNA–protein; n = 110), OSMR (0.340; 0.410; n = 89), and LHFPL2 (0.261; 0.217, P = 0.061; n = 75). These associations provide protein-level support in the analyzed HNSCC cohort, strongest for MICAL2, whereas the LHFPL2 mRNA– protein correlation did not reach nominal significance.

**Figure 10.**
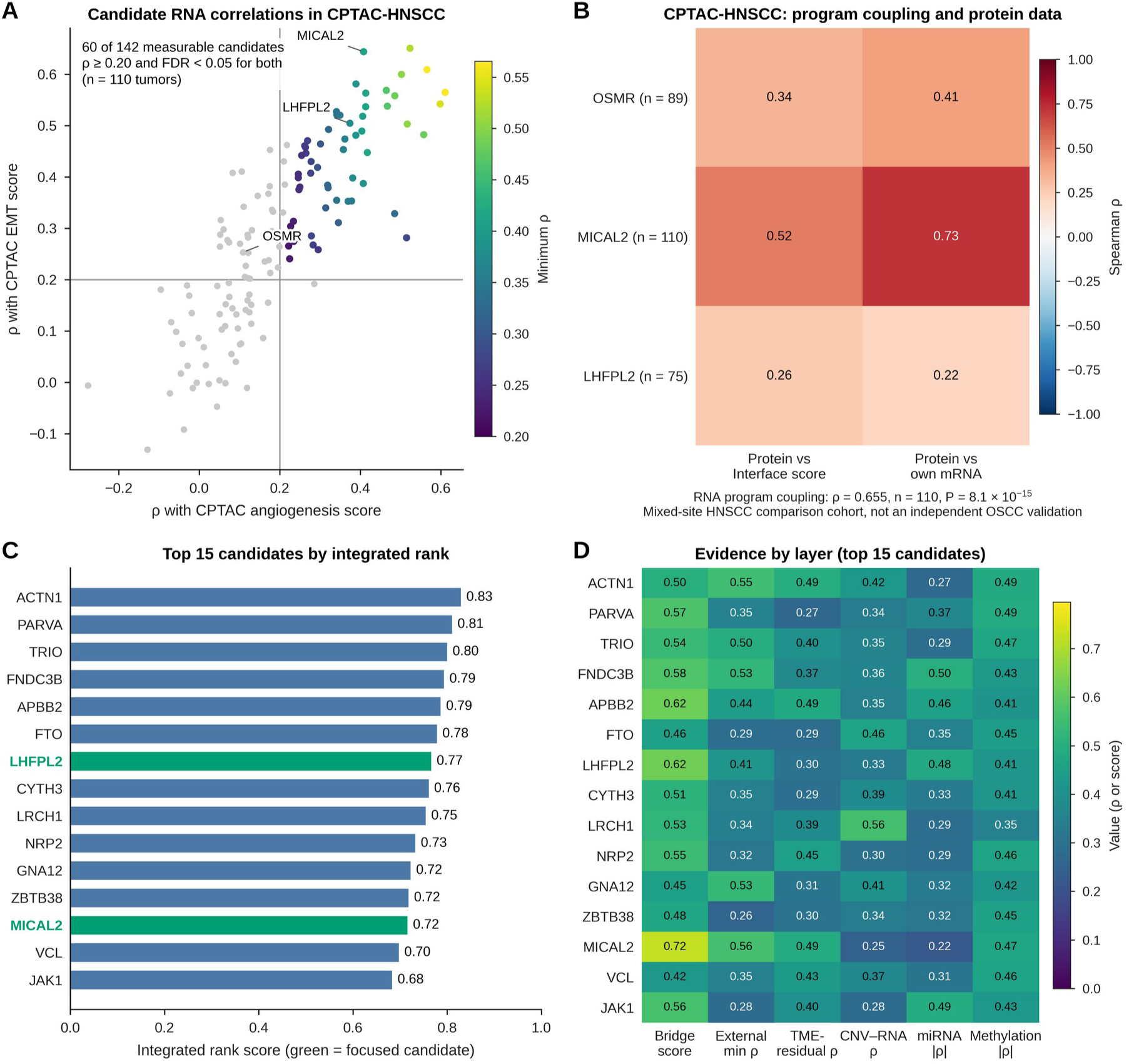
CPTAC-HNSCC analysis and integrated ranking of the final candidates. (A) RNA correlations of the 142 measurable final candidates with the CPTAC angiogenesis and EMT scores in 110 tumors; colored points met ρ ≥ 0.20 and FDR < 0.05 for both. (B) Spearman correlations of protein abundance with the RNA-based Interface score and with the gene’s own mRNA for the focused candidates (n, tumors with protein values). CPTAC-HNSCC is a mixed-site HNSCC comparison cohort, not an independent OSCC validation cohort. (C) The 15 final candidates with the highest integrated rank scores; focused candidates are shown in green. (D) Evidence-layer values for the same candidates.

In addition, we ranked the final candidates by an integrated rank score derived from the combined evidence matrix. Fig. 10C shows the 15 highest-ranked candidates, led by ACTN1, PARVA, TRIO, FNDC3B, and APBB2; among the focused candidates, LHFPL2 ranked seventh and MICAL2 thirteenth, whereas OSMR was not among the top 15.

Finally, we evaluated which specific combinations of evidence support the individual high-ranked candidate genes. Fig. 10D shows that the top-ranked candidate genes were supported by diverse and complementary combinations involving discovery strength, external-cohort support, associations retained after TME adjustment, CNV–RNA associations, predicted miRNA targeting, and promoter methylation, indicating different combinations of supporting evidence across candidates.

### Signature robustness and clinical associations of the angiogenesis–EMT state

Next, we checked whether program coupling depends on the custom gene sets by replacing them with the standardized MSigDB Hallmark gene sets. Fig. 11A shows strong positive program coupling across all signature combinations: Hallmark angiogenesis with custom EMT (ρ = 0.785), custom angiogenesis with Hallmark EMT (ρ = 0.816), and both Hallmark signatures (ρ = 0.889), supporting robustness of the association across the tested signature combinations. Because the Hallmark angiogenesis and EMT gene sets share 12 genes (12 of the 36 Hallmark angiogenesis genes), the Hallmark–Hallmark coupling may partly reflect shared genes.

**Figure 11.**
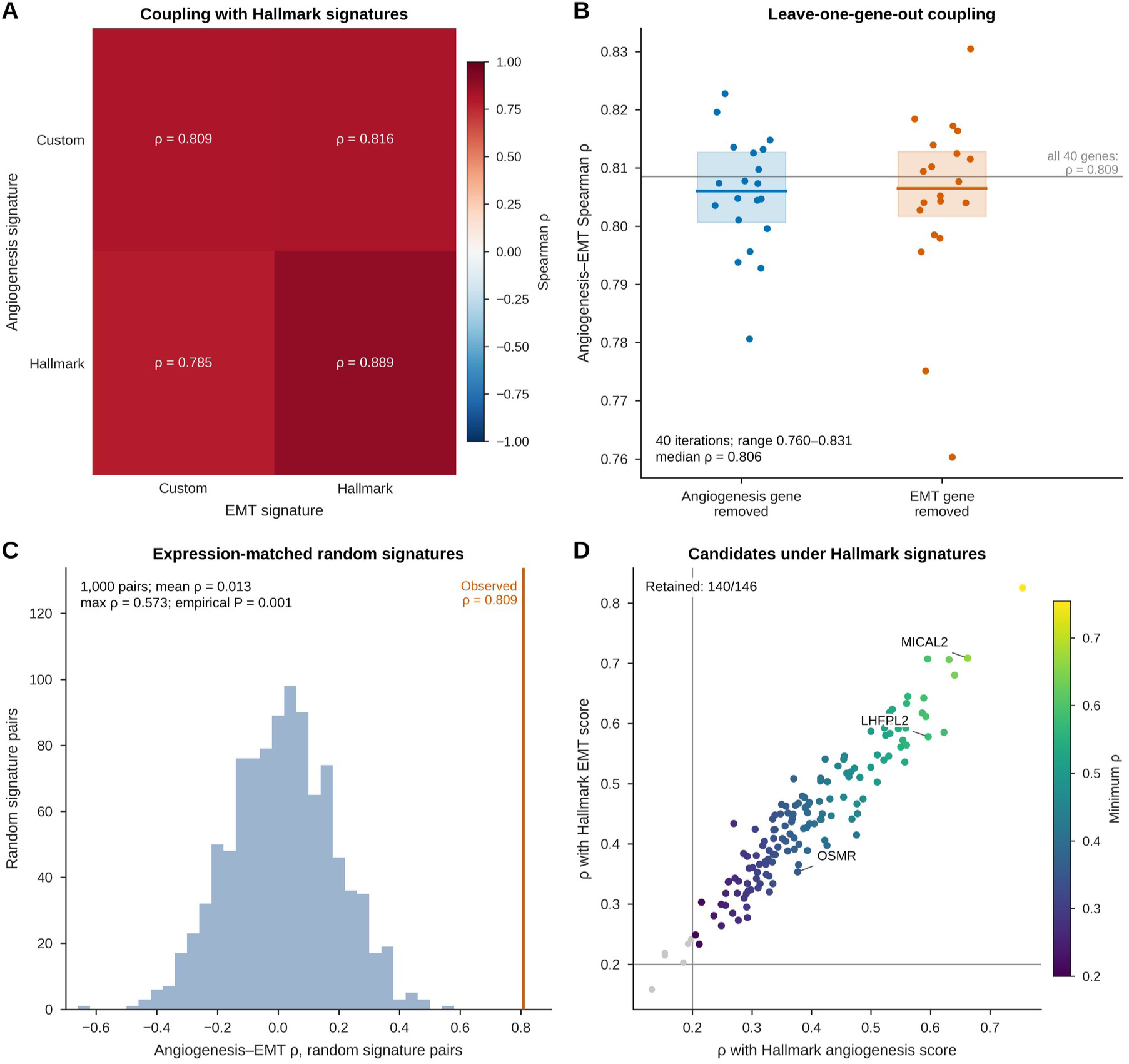
Robustness of the angiogenesis–EMT coupling. (A) Coupling with the custom and MSigDB Hallmark signatures. (B) Coupling after leaving out each of the 40 signature genes in turn; boxes show interquartile ranges and lines medians, and the gray line marks the value with all 40 genes. (C) Coupling for 1,000 expression-matched random signature pairs; the orange line marks the observed value. (D) Correlations of the final candidates with the Hallmark-based angiogenesis and EMT scores, colored by the smaller ρ; lines mark ρ = 0.20, and gray points were not retained.

Additionally, we tested whether this program coupling depends on a single constituent gene by removing one gene at a time from the program signatures. Fig. 11B shows that, across 40 leave-one-gene-out iterations, the correlation remained strongly positive (ρ range, 0.760–0.831; median ρ = 0.806). The positive association therefore persisted after removal of each individual signature gene.

Further, we compared the angiogenesis–EMT correlation against an empirical null distribution of 1,000 pairs of expression-matched random gene signatures. Fig. 11C shows that the null distribution yielded a mean ρ of 0.013, with no random signature pair attaining the observed value of 0.809 (maximum null ρ = 0.573; empirical P = 0.001), indicating that the observed correlation exceeded those generated under the tested null model.

Finally, we calculated the proportion of final candidates that retain dual positive program coupling under the standardized Hallmark signatures. Fig. 11D shows that 140 of 146 final candidates retained a dual-program association under the MSigDB Hallmark scoring framework, indicating the robustness of the candidate bridge-gene support.

We further examined the relationship between the Interface score, clinicopathological characteristics, and overall survival (OS) in OSCC. Interface scores overlapped across pathological stages, with no clear increase from early to advanced stages (Kruskal–Wallis test; n = 310; P = 0.339; Fig. 12A). Scores also showed overlapping distributions across nodal-status categories, without clear separation between node-positive and node-negative tumors (Mann– Whitney test; n = 286; P = 0.214; Fig. 12B). Kaplan–Meier analysis detected no statistically significant difference in OS between the low-and high-Interface-score groups (n = 170 each; log-rank P = 0.21; Fig. 12C). No statistically significant association between the continuous Interface score and OS was detected in the univariable (HR per 1-SD increase = 0.92, 95% CI 0.78–1.08; P = 0.295; n = 340, 154 events), stage-adjusted (HR = 0.92, 95% CI 0.78–1.08; P = 0.318; n = 308, 141 events), or nodal-adjusted (HR = 0.93, 95% CI 0.78–1.09; P = 0.367; n = 284, 124 events) analyses (Fig. 12D). Proportional-hazards diagnostics showed no notable violations of the survival-model assumptions evaluated (all P > 0.05; Fig. 13A).

**Figure 12.**
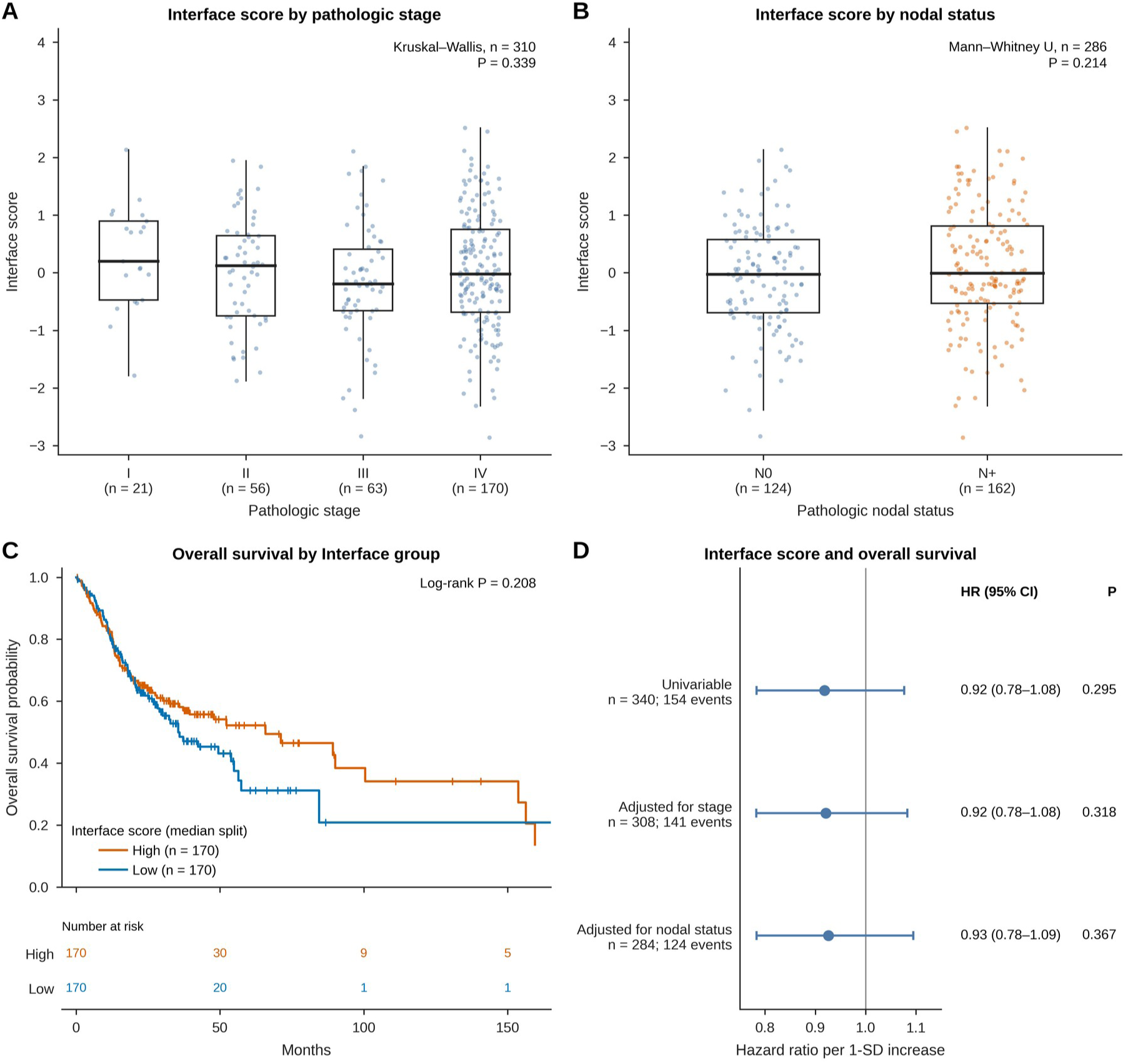
Clinicopathological and survival associations of the Interface score. (A) Interface score by pathologic stage (Kruskal–Wallis test). (B) Interface score by pathologic nodal status (two-sided Mann–Whitney U test). (C) Kaplan–Meier curves of overall survival for tumors split at the median Interface score (log-rank test), with numbers at risk. (D) Hazard ratios per 1-SD increase in the Interface score with 95% confidence intervals from univariable, stage-adjusted and nodal-adjusted Cox models.

**Figure 13.**
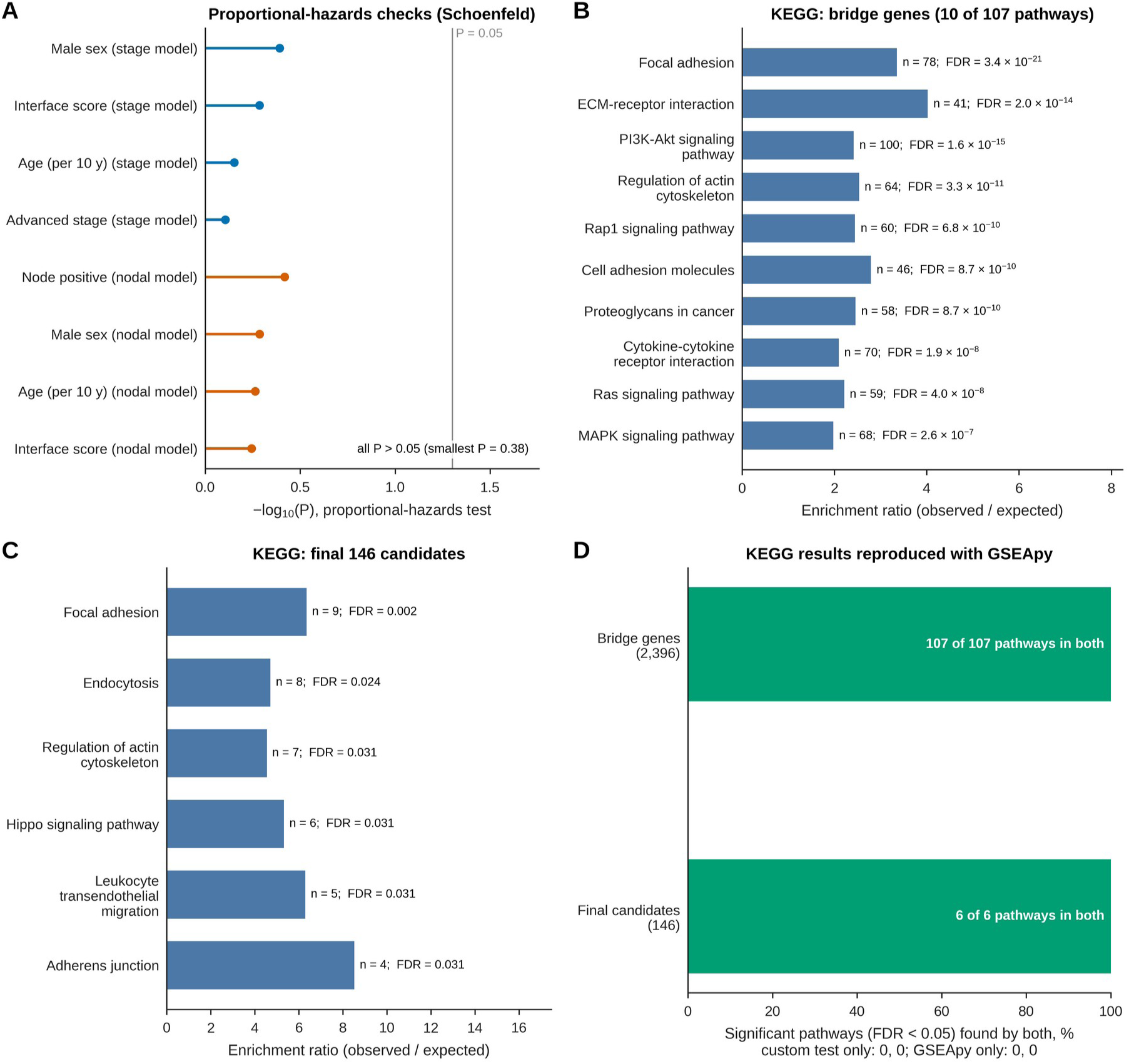
Proportional-hazards checks and KEGG pathway enrichment. (A) −log10(P) of Schoenfeld-residual tests for the covariates of the stage-adjusted and nodal-adjusted Cox models; the gray line marks P = 0.05. (B) Ten of the 107 KEGG pathways significantly enriched among the bridge genes, with the number of overlapping genes and FDR. (C) The six KEGG pathways significantly enriched among the final candidates. (D) Agreement between the custom Fisher’s exact tests and GSEApy in the significant pathways (FDR < 0.05).

### Pathway-level analyses place the bridge-gene architecture within coordinated adhesion, matrix, and signaling programs

Finally, we assessed which biological pathways were enriched among the bridge genes and among the final candidates. Of the 107 KEGG pathways significantly enriched in the larger bridge-gene collection, Fig. 13B shows 10 selected pathways, led by focal adhesion (FDR = 3.4 × 10^−21^), extracellular matrix–receptor interaction, PI3K–Akt signaling, regulation of actin cytoskeleton, Rap1 signaling, cell adhesion molecules, proteoglycans in cancer, cytokine– cytokine receptor interaction, Ras signaling, and MAPK signaling.

Six KEGG pathways were significantly enriched among the 146 final candidates (Fig. 13C): focal adhesion (FDR = 0.002), endocytosis (FDR = 0.024), and regulation of actin cytoskeleton, Hippo signaling, leukocyte transendothelial migration, and adherens junction (each FDR = 0.031). Reanalysis with GSEApy reproduced all 107 significant pathways for the bridge genes and all six for the final candidates, with no pathway significant in only one implementation (Fig. 13D).

## Discussion

Angiogenesis and EMT are two of the key hallmarks of oral squamous cell carcinoma (OSCC) that are frequently driven by shared microenvironmental cues associated with fibrovascularization and ECM remodeling (Wei et al., 2015; Bordeleau et al., 2017; Pomella et al., 2024). Although both are triggered by common cues, few studies have shown the effect of one program on the other. For example, Takahashi et al. (2020) reported that inhibiting TGF-β signaling blocked TGF-β-induced EMT in oral cancer cells and reduced angiogenesis in an OSCC xenograft model. Despite these shared stimuli (Pomella et al., 2024), it remains unclear whether these two major hallmark processes are coordinated as whole transcriptional programs across OSCC tumors, rather than linked only through individual factors. Therefore, in this paper, we used different approaches to assess whether angiogenesis and EMT are coordinated across OSCC tumors via transcriptome-wide, composition-adjusted, multi-omic, and external-cohort analyses. Our analysis highlighted a strong correlation between angiogenesis and EMT across 342 TCGA-OSCC tumors (Spearman ρ = 0.809; Fig. 1A), and this coupling extended to 2,396 transcriptome-wide bridge genes (Fig. 1B). In addition, this coupling was robust to alternative signature definitions (Fig. 11A,B) and was reproduced in an external OSCC cohort (GSE30784; ρ = 0.648; Fig. 9A). This suggests that angiogenesis and EMT are not independent features but may represent components of a broader remodeling state, in which vascular and EMT-related transcriptional programs rise together. In this interpretation, vascular remodeling supports growth (Carmeliet & Jain, 2011), whereas epithelial plasticity supports motility and invasion (Kalluri & Weinberg, 2009), and both programs support OSCC progression (Pomella et al., 2024). In this context, such a coordinated state is plausible: OSCC is a heterogeneous cancer type (The Cancer Genome Atlas Network, 2015; Leemans et al., 2018) in which shared AKT and MAPK signaling can promote both EMT and new vessel formation (Pomella et al., 2024). Part of this coupling, however, reflects tumor composition (ρ = 0.406 after adjustment for purity and stromal and immune content; Fig. 3C), as discussed below.

To examine in depth the role of bridge genes in the coupling of angiogenesis and EMT, we further analyzed the functional pathways and structural components that are common across these bridge genes. The characterization of the bridge genes revealed that they are associated with a common fibrovascular and ECM microenvironment, which connects features of both EMT and angiogenesis. On closer analysis, we found that the top bridge genes positively correlated with both programs included LAMA4 and NID2 (Fig. 1C) (maturation and integrity of the endothelial basement membrane) (Thyboll et al., 2002; Bader et al., 2005), ITGA1 (collagen adhesion and tumor vascularization) (Pozzi et al., 2000; Ghatak et al., 2016), PDGFRB (recruitment of pericytes) (Hellström et al., 1999; Sugimoto et al., 2006), and ADGRA2 (GPR124) (endothelial sprout development) (Carson-Walter et al., 2001; Kuhnert et al., 2010). The functional annotation of the bridge genes also supports this ECM remodeling context (Fig. 2C), and the 107 significantly enriched KEGG pathways were mainly led by focal adhesion, ECM–receptor interaction, PI3K–Akt signaling, regulation of actin cytoskeleton and MAPK signaling (Fig. 13B). This suggests a possible mechanism through which EMT and angiogenesis could be coupled. Experiments suggest that ECM remodeling and stiffening of the matrix trigger the clustering of integrins, which therefore promotes focal adhesion assembly and downstream activation of PI3K–Akt signaling (Paszek et al., 2005; Levental et al., 2009), eventually leading to cell migration and mesenchymal transition (Wei et al., 2015), including in OSCC cells (Matte et al., 2019), while simultaneously driving endothelial sprouting and vascular permeability (Bordeleau et al., 2017). Although hypoxia can drive both programs via HIF-dependent pathways (Pomella et al., 2024), the hypoxia gene set was not significantly enriched (FDR = 0.531; Fig. 2C), within the limits of bulk hypoxia signatures. This underscores that our finding reflects a stromal/ECM-associated transcriptional state, where matrix–integrin signaling may provide a shared microenvironment for dual-program co-variation, although neither matrix stiffness nor pathway activation was measured here. However, this stromal/ECM-associated state involves both malignant and non-malignant cells, and higher stromal content alone could produce a similar pattern. This led us to determine whether the coupling is retained after statistically adjusting for stromal and endothelial cell composition, and whether EMT features also exist in malignant cells.

Single-cell expression analyses of the final candidate genes point to angiogenesis–EMT coupling residing in several tumor compartments. Our results also suggest that, of the 146 final candidates, 41 have predominant expression in endothelial cells, followed by fibroblasts (26), malignant cells (20), and macrophages (18) (Fig. 4A). Additionally, there is a strong association between CAF marker scores and the angiogenesis, EMT, and Interface scores (ρ = 0.80–0.84), while endothelial marker scores are most strongly associated with angiogenesis (ρ = 0.82) and the Interface (ρ = 0.73), as shown in Fig. 3A. After we adjusted for tumor purity and for stromal and immune infiltration, the angiogenesis–EMT association decreased but remained positive (ρ = 0.406; Fig. 3C). This pattern shows a greater reduction with stromal/immune adjustment (ρ = 0.400) than with tumor purity adjustment alone (ρ = 0.695). This suggests that CAF and endothelial compartments contribute to the Interface, possibly through paracrine signaling between fibroblasts and neighboring malignant cells, which may further influence vascular remodeling. Looking at known mechanisms, CAFs support ECM deposition (Kalluri, 2016; Sahai et al., 2020) and secrete growth factors and chemokines that bolster angiogenesis by recruiting pericytes and endothelial cells (Kalluri & Zeisberg, 2006). Further, CAFs secrete TGF-β1, which supports the invasion of OSCC cells, with changes in EMT markers (Haga et al., 2021). Thus, CAFs are a plausible source for driving both programs simultaneously. Additionally, tumor endothelial cells are heterogeneous and, beyond angiogenesis, are linked to immune regulation and matrix organization (Zeng et al., 2023). Further, when we scored EMT with an epithelial-restricted gene set, the coupling remained strong (ρ = 0.710; Fig. 4C); the Interface score also showed an association with pEMT (ρ = 0.59) and retained the same pattern after purity adjustment (Fig. 5B). After removing the two genes shared by the pEMT and EMT signatures (MMP2 and VIM), the association was still retained (ρ = 0.554), pointing to a possible involvement of partial, intermediate EMT states (Nieto et al., 2016) rather than stromal contamination alone. However, these bulk scores cannot show that this signal arises within malignant cells. A prior single-cell study has reported a biological basis for this observation, in which malignant cells expressing pEMT programs are localized near the invasive edge of oral cavity tumors, and has identified potential ligand–receptor interactions, such as TGFB3– TGFBR2, FGF7–FGFR2, and CXCL12–CXCR7, which suggest plausible routes through which stromal cells influence malignant cell plasticity (Puram et al., 2017). Taken together, the data suggest a working model in which shared microenvironmental signals could trigger both epithelial plasticity and vascular remodeling simultaneously.

Additionally, we performed multi-omic analyses, which revealed genomic and epigenomic associations that may help explain why candidate bridge-gene expression varies across tumors. Copy number was positively associated with expression in a subset of bridge genes (Fig. 6A), suggesting that localized gene amplification may contribute to their elevated expression. From an epigenetic point of view, candidate gene expression was inversely associated with promoter methylation (Fig. 6C). This inverse relationship was largely limited to the promoter region: genome-wide CpG sites showed mixed, mostly positive associations with the Interface (Fig. 7A), pointing to the need for site-specific functional annotation. Moreover, candidate bridge-gene expression and the Interface were inversely associated with microRNAs, including the miR-200 family and miR-205 (Fig. 7B–D), which suppress EMT by directly targeting the transcriptional repressors ZEB1 and SIP1 (ZEB2), as shown in non-OSCC epithelial cells (Gregory et al., 2008). As miR-200 also inhibits angiogenesis in other cancers (Pecot et al., 2013), loss of these miRNAs could derepress targets in both programs, consistent with the coupling we observed. Most of our miRNA–gene pairs, however, were predicted rather than experimentally established, and only a small subset had strong experimental support in miRTarBase (Fig. 8A; Cui et al., 2025). Together, copy-number gain, promoter hypomethylation, and miRNA loss may contribute to the expression of the bridge genes, providing a rationale for evaluating prioritized candidates such as OSMR, MICAL2, and LHFPL2 (Fig. 6D). Because copy number and methylation also guided candidate selection, their support for these candidates is not independent. In addition, the methylation and miRNA analyses were not adjusted for tumor purity, which affects both bulk methylation (Zheng et al., 2017) and miR-200, an epithelial marker (Park et al., 2008). These associations are therefore hypothesis-generating rather than evidence of regulation. Nonetheless, each of these three candidates is associated with both the angiogenesis and EMT programs. Mechanistically, OSMR encodes a cell-surface receptor for the inflammatory cytokine oncostatin M (OSM; Caffarel & Coleman, 2014). In cervical squamous cell carcinoma cells, OSMR overexpression promoted OSM-induced, STAT-dependent VEGF-A expression, endothelial tube formation, migration, and invasion (Winder et al., 2011), and was associated with EMT (Kucia-Tran et al., 2016). In OSCC, high OSMR expression has been associated with JAK/STAT signaling, lymph node metastasis, and poorer survival (Geng et al., 2023). In pancreatic cancer, macrophage-derived OSM reprogrammed fibroblasts toward an inflammatory, pro-tumorigenic state (Lee et al., 2021), which fits the fibroblast-dominant OSMR expression in our data (Fig. 4B). MICAL2 regulates actin filament disassembly (Lundquist et al., 2014) and promotes invasive cell motility (Mariotti et al., 2016); it is also expressed in tumor neo-angiogenic endothelium, where it is required for the endothelial response to VEGF (Barravecchia et al., 2019). In contrast, the mechanism of action of LHFPL2 is less well defined. One computational study linked LHFPL2 expression to macrophage infiltration and M2 polarization in renal cell carcinoma (Gong et al., 2024), consistent with its macrophage-dominant expression in our data (Fig. 4B), but no direct role in coordinating both programs has been shown. Our study also revealed RNA–protein concordance and protein–Interface associations in CPTAC-HNSCC (Fig. 10B), which further support the expression patterns of OSMR and MICAL2, whereas protein support for LHFPL2 was weaker. Overall, these observations suggest that the candidate genes may influence distinct aspects of the shared tumor state, including cellular plasticity and cytokine responsiveness, rather than acting through a single pathway.

Across our analyses, angiogenesis–EMT coordination was retained in different datasets and external cohorts, suggesting that it is a stable biological feature rather than a dataset-specific artifact. In the external OSCC cohort (GSE30784; Chen et al., 2008), the dual-program scores maintained a strong positive correlation (Fig. 9A), and the coupling also recurred in CPTAC-HNSCC (Fig. 10B; Huang et al., 2021) and across pan-cancer datasets (Fig. 9D). Its recurrence in 12 cancer cohorts, including glioblastoma, also shows that the coupling is not specific to OSCC. This cross-cohort reproducibility is consistent with a recurrent multicellular state in which pEMT carcinoma cells may exchange paracrine cues with endothelial cells and CAFs during ECM remodeling (Carmeliet & Jain, 2011; Puram et al., 2017). However, it does not test this model, and recurring stromal content could also contribute. Sensitivity analyses also showed that the coordination was resilient to signature substitution (Fig. 11A) and leave-one-gene-out analysis (Fig. 11B), and far exceeded an expression-matched random-gene null (Fig. 11C). However, replication of the overall coupling should be distinguished from replication of the candidates: GSE30784 was also used to select the candidates (Fig. 9C), so CPTAC-HNSCC provided their only independent test, in which only a subset replicated (Fig. 10A) and proteomic data supported some of them (Fig. 10B). Thus, the dual-program coupling is robust and reproducible at the transcript level, whereas the candidate-level findings require further independent validation.

Our results also indicate that this coordination of angiogenesis and EMT does not point towards prognostic utility, as clinical evaluation revealed that Interface score distributions overlapped across different pathological stages (Kruskal–Wallis P = 0.339; Fig. 12A) and nodal categories (Mann–Whitney P = 0.214; Fig. 12B), indicating that program magnitude may not increase with disease progression. The Interface also showed no significant association with overall survival (log-rank P = 0.21; Cox HR 0.92 per SD, 95% CI 0.78–1.08, with similar estimates after adjustment for stage or nodal status; Fig. 12C, D), although a nonsignificant result does not demonstrate the absence of an effect (Altman & Bland, 1995), which may indicate continuous paracrine crosstalk between pEMT carcinoma cells, sprouting vessels, and stroma throughout progression (Puram et al., 2017). However, bulk tissue profiling mixes cell types, and CPTAC proteomics include non-OSCC subsites, with only 49 of 110 tumors from the oral cavity. These limitations, along with the lack of spatial mapping and functional experiments, mean this study is a prioritized candidate-gene framework rather than a validated clinical marker (Aran et al., 2015; Huang et al., 2021; Arora et al., 2023).

Thus, this provides multi-cohort evidence of a transcriptomic association, and we propose a biphasic invasive model in which ECM and cell–matrix interactions stimulate transcriptional responses and influence bridge gene expression. Ultimately, this is a framework for systematically testing how microvascular expansion and epithelial plasticity functionally interact in the OSCC microenvironment. In the future, high-resolution spatial transcriptomics (Simkin et al., 2026) and cell-type-specific approaches in 3D microfluidic co-cultures are required to validate these targets.

## Data and code availability

TCGA HNSC RNA-seq (TCGA_HNSC_HiSeqV2), GISTIC2 copy-number, HumanMethylation450 methylation, mature miRNA and clinical data are available from UCSC Xena (https://xenabrowser.net). GSE30784, GSE103322, GSE37991 and GSE41613 are available from the Gene Expression Omnibus (https://www.ncbi.nlm.nih.gov/geo/). CPTAC-HNSCC data are available from the cptac Python package (version 1.5.14; https://github.com/PayneLab/cptac; Lindgren et al., 2021). Gene sets were obtained from MSigDB (Hallmark collection, v2023.1) and the Enrichr KEGG_2021_Human library, and miRNA–target interactions from miRTarBase release 10.0. The analysis code and the processed tables underlying each figure are available from the corresponding author on request and will be deposited in a public repository.

## Ethics statement

This study used only publicly available, de-identified data, and no new human samples or data were collected. Institutional review board approval was therefore not required.

## Acknowledgements

This research was supported by the Smathers Libraries HiPerGator Sponsorship Program (https://guides.uflib.ufl.edu/HiPerGatorSponsorship). The authors acknowledge UFIT Research Computing for providing computation resources and support that have contributed to the research results reported in this publication (https://it.ufl.edu/rc).

## Funding

This research was supported by the Smathers Libraries HiPerGator Sponsorship Program, University of Florida.

## Author contributions

S.D.: Conceptualization, Data curation, Formal analysis, Funding acquisition, Investigation, Methodology, Project administration, Resources, Software, Supervision, Validation, Visualization, Writing – original draft, Writing – review and editing. E.N.: Conceptualization. R.Si.: Data curation, Writing – review and editing. R.Sh.: Data curation, Writing – review and editing. All authors read and approved the final manuscript.

## Competing interests

The authors declare no competing interests.

## Supplementary information

**Supplementary Table S1.** Correlation thresholds, FDR criteria and multiple-testing families used in each analysis.

| Step | Correlation threshold | FDR | Correction family |
|---|---|---|---|
| Bridge genes (TCGA-OSCC) | ρ ≥ 0.30 with both scores | < 0.05 for both | 20,213 genes, per program |
| TME-adjusted Interface support | Absolute ρ ≥ 0.20 | < 0.05 | 2,396 bridge genes |
| Program-specific TME support | ρ ≥ 0.20 with both residual scores | < 0.05 for both | Bridge genes, per program |
| Copy-number support | ρ > 0.20 | < 0.05 | 2,274 tested genes |
| Methylation support | Absolute ρ ≥ 0.20 | < 0.05 | 396,065 CpGs |
| Promoter methylation | ρ ≤ −0.20 with the Interface and with expression | < 0.05 for both | CpGs (Interface); 303 CpG–gene pairs (expression) |
| High-confidence miRNAs | Absolute ρ ≥ 0.20 | < 0.05 | 1,101 miRNAs |
| Inverse miRNA–target pairs | ρ ≤ −0.20 | < 0.05 | 3,422 pairs |
| GSE30784 replication | ρ ≥ 0.20 with both scores (stricter subset ≥ 0.30) | < 0.05 for both | 302 mapped candidates, per program |
| Composition-adjusted candidates (ESTIMATE, cell fractions) | ρ ≥ 0.20 with both scores | < 0.05 for both | 146 candidates, per program |
| CPTAC and Hallmark persistence | ρ ≥ 0.20 with both scores | < 0.05 for both | 142 or 146 candidates, per program |
| Bootstrap stability | ρ ≥ 0.30 with both scores | Unadjusted P < 0.05 | Not corrected |

## Notes

https://www.ncbi.nlm.nih.gov/geo/query/acc.cgi?acc=GSE30784

https://www.ncbi.nlm.nih.gov/geo/query/acc.cgi?acc=GSE103322

https://www.ncbi.nlm.nih.gov/geo/query/acc.cgi?acc=GSE37991

https://www.ncbi.nlm.nih.gov/geo/query/acc.cgi?acc=GSE41613

https://xenabrowser.net

https://github.com/PayneLab/cptac

### Competing Interest Statement

The authors have declared no competing interest.

